# Chitosan biosynthesis and virulence in the human fungal pathogen *Cryptococcus gattii*

**DOI:** 10.1101/759050

**Authors:** Woei C. Lam, Rajendra Upadhya, Charles A. Specht, Abigail E. Ragsdale, Camaron R Hole, Stuart M. Levitz, Jennifer K. Lodge

## Abstract

*Cryptococcus gattii* R265 is a hyper-virulent fungal strain responsible for the major outbreak of cryptococcosis in Vancouver Island of British Columbia in 1999. It differs significantly from *C. neoformans* in its natural environment, its preferred site in the mammalian host, and in the nature and mode of pathogenesis. Our previous studies in *C. neoformans* have shown that the presence of chitosan, the deacetylated form of chitin, in the cell wall attenuates inflammatory responses in the host, while its absence induces robust immune responses, which in turn facilitate clearance of the fungus and induces a protective response. The results of the present investigation reveal that the cell wall of *C. gattii* R265 contains 2-3-fold higher amount of chitosan compared to that of *C. neoformans*. The genes responsible for the biosynthesis of chitosan are highly conserved in the R265 genome; the roles of the three chitin deacetylases (CDA) have however, been modified. To deduce their roles, single, double and a triple *CDA* deletion strains were constructed in a R265 background and were subjected to mammalian infection studies. Unlike *C. neoformans* where Cda1 has a discernible role in fungal pathogenesis, in R265 Cda3 is critical for virulence. Deletion of either *CDA3* alone (*cda3Δ*) or in combination with either *CDA1* (*cda1Δ3Δ*) or *CDA2* (*cda2Δ3Δ*) or both (*cda1Δ2Δ3Δ*) rendered the yeast cells avirulent and were cleared from the infected host. Moreover, the *cda1Δ2Δ3Δ* strain of R265 induced a protective response to a subsequent infection with R265. These studies shed more light into the regulation of chitosan biosynthesis of *C. gattii* and its subsequent effect on fungal virulence.

**Importance:** The fungal cell wall is an essential organelle whose components provide the first line of defense against host-induced antifungal activity. Chitosan is one of the carbohydrate polymers in the cell wall that significantly affects the outcome of host-pathogen interaction. Chitosan-deficient strains are avirulent, implicating chitosan as a critical virulence factor. *C. gattii* R265 is an important fungal pathogen of concern due to its ability to cause infections in individuals with no apparent immune dysfunction and an increasing geographical distribution. Characterization of the fungal cell wall and understanding the contribution of individual molecules of the cell wall matrix to fungal pathogenesis offers new therapeutic avenues for intervention. In this report, we show that the *C. gattii* R265 strain has evolved alternate regulation of chitosan biosynthesis under both laboratory growth conditions and during mammalian infection compared to that of *C. neoformans*.

## Introduction

Cryptococcosis is an invasive fungal infection caused mainly by the Cryptococcus species *C. neoformans* and *C. gattii*. Cryptococcus is ubiquitous with world-wide distribution and causes infections in a wide variety of host species, such as plants, birds and mammals (1, 2). Infections caused by Cryptococcus may lead to cryptococcal meningitis and is estimated to cause more than 200,000 deaths annually (3). *C. neoformans* (inclusive of serotypes A and D) is an opportunistic pathogen and mainly causes disease in immune-compromised patients (4, 5). However, infections even in healthy individuals (non-HIV) with or without underlying risk factors, have also been reported (6, 7). Unlike *C. neoformans*, its sibling species *C. gattii* (serotypes B and C) is recognized as a primary pathogen as it predominately causes infections in immune competent individuals (5, 8–10). *C. gattii* was initially considered to be endemic to tropical and subtropical regions, especially Australia until it attracted attention with a major outbreak on Vancouver Island, British Columbia, in 1999 (11). Based on the global molecular epidemiologic survey employing a wide variety of molecular techniques, five distinct genetic groups (VGI/AFLP4, VGII/AFLP6, VGIII/AFLP5, VGIV/AFLP7 and VGIV/AFLP10) within the *C. gattii* species complex were described (12). The strain *C. gattii* R265 belongs to the VGII subtype and was the strain associated with the outbreak in British Columbia in 1999 (13). The fungus subsequently spread to the Pacific Northwest of the United States (14–16). Infections due to *C. gattii* R265 are associated with an 8-20% mortality rate in spite of anti-fungal therapies. With advances in genotyping, *C. gattii* distribution has been revised: It is predominantly isolated from environmental sources and is currently found in a variety of climates, including humid and arid conditions and associated with 53 different tree species across six continents (10). It has also been isolated from diverse groups of organisms, including cats, dogs, marine mammals, koalas, deer, ferrets, llamas, horses, birds and insects (10). This distribution to diverse conditions of environment and host species demonstrates its adaptability.

Significant differences in the ecological, morphological, biochemical, molecular, pathological and clinical features exist between *C. neoformans* and *C. gattii* species complex (12, 17). For example, the difference in the assimilation of nitrogen and carbon sources between *C. neoformans* and *C. gattii* species has been exploited to formulate a one-step diagnostic media for their identification (18). Several differences in the nature of the host-immune response elicited by *C. gattii* R265 compared to *C. neoformans* has been attributed to its capacity to cause disease in persons with apparently normal immune system. *C. gattii* R265 has been shown to proliferate better in the macrophages compared to *C. neoformans* due to their increased resistance to reactive oxygen species (ROS) inside macrophages and to their tubular mitochondrial morphology (19). At the genomic level, R265 has shown to evolve by expanding stress related heat shock proteins, which may offer better fitness inside the host (20). *C. gattii* R265 readily proliferates in the lung and disseminates poorly to the brain, suggesting that unlike *C. neoformans*, the major target organ of *C. gattii* R265 is the host lung (19, 21). *C. gattii* R265 has been shown to trigger a dampened immune response in the lung with reduced infiltration of macrophages, neutrophils and Th1/Th17 lymphocytes, and it inhibits host dendritic cell maturity (22–25). Both *C. neoformans* and *C. gattii* R265 share a similar suite of virulence factors, yet they differ in the nature of pathogenesis in mammalian hosts.

Chitosan is one of the critical components of *C. neoformans* cell wall and shown to be essential for its virulence (26). The genes coding for enzymes that are responsible for the production of chitosan in *C. neoformans* have been identified and characterized (27). Out of the eight potential chitin synthase genes in the genome, chitin synthase 3 (Chs3) coded by *CHS3* and chitin synthase regulator 2 (Csr2) coded by the *CSR2* gene are critical for the production of chitosan (27). There are four potential chitin deacetylase (*CDA*) genes in the genome of *C. neoformans*; three have been shown to possess chitin deacetylase activity in vegetatively growing cells (28). Chitosan-deficient strains of *C. neoformans* induce significant host-immune responses during mammalian infection, and clearance of the fungus (26, 29). These results suggest that presence of chitosan may influence the cell wall architecture, thereby shielding pathogen associated molecular patterns (PAMPSs) from being recognized by host immune cells. When grown in YPD culture medium, all three Cda proteins appear to be functionally redundant. However, they are differentially regulated in the lungs of the infected host with Cda1 being preferentially expressed (30). Accordingly, either *cda1Δ* or the *cda* mutant with abolished chitin deacetylase activity were found to be avirulent suggesting that Cda1 alone with its chitin deacetylase activity is sufficient to render the yeast cells fully virulent during a murine infection of CBA/J mice. Deletion of either Cda2 or Cda3 did not affect the virulence of the yeast strains (30). These results indicate that in *C. neoformans*, chitosan and the mechanisms of its production are the critical mediators of fungal pathogenesis and virulence.

The mechanisms responsible for differences in the pathogenicity and the virulence composite between *C. neoformans* and *C. gattii* are not understood. We sought to determine if there are any differences in either the level of cell wall chitosan or in the regulation of its biosynthesis between these two species. Here, we report the identification of genes present in *C. gattii* R265 genome that are either responsible for the production of chitosan during growth under vegetative conditions or that contribute to chitosan biosynthesis during mammalian infection. We found significant differences in the amount and the regulation of chitosan biosynthesis between *C. gattii* R265 and *C. neoformans*. First, the cell walls of *C. gattii* R265 had more than double the amount of chitosan compared to *C. neoformans* when either grown in culture or in infected mice. We targeted homologs of *C. neoformans* chitosan biosynthetic genes in the *C. gattii* R265 genome for gene deletion and subjected the respective deletion strains to various *in vitro* phenotypic assays and to mouse virulence studies. For *C. neoformans*, Cda1 played an important role in the synthesis of chitosan, while Cda3 was found to be dispensable during murine infection. Interestingly for *C. gattii* R265, we found that Cda3 plays a critical role in fungal virulence, while the deletion of Cda1 did not affect fungal virulence across different mouse strains. The results of these studies will provide a framework to further design strategies to dissect the molecular mechanisms of chitosan in fungal induced host-immune response and virulence.

## Results

### Identification of *C. gattii* R265 genes potentially involved in the synthesis of chitosan

In *C. neoformans*, the conversion of chitin to chitosan is catalyzed by Cda1, Cda2 and Cda3 (28, 30). We utilized BLASTp homology with *C. neoformans* Cda1, Cda2, and Cda3 to identify the *C. gattii* chitin deacetylases in the R265 genome. This search yielded CNBG_1745 (Cda1), CNBG_9064 (Cda2) and CNBG_0806 (Cda3) as the *C. gattii* homologs of *C. neoformans* (Table 1). At the protein level, Cda1, Cda2 and Cda3 are 85%, 83% and 85% identical and 92%, 90% and 90% similar, respectively, between the two species. All three Cda proteins of *C. gattii* share similar predicted sequence features to that of *C. neoformans* Cda proteins in having N-terminal signal sequences, S/T rich regions and GPI anchor sites, as well as conserved amino acids for catalysis. Pairwise protein sequence alignments are shown in Supplementary Fig. 1.

**Table 1.** Identification of chitin deacetylases (CDA) in *C. gattii* R265 genome.

| Locus tag | *C. gattii*<br>protein | E value for<br>*C. neoformans*<br>Cda1p query | E value for<br>*C. neoformans*<br>Cda2p query | E value for<br>*C. neoformans*<br>Cda3p query |
| --- | --- | --- | --- | --- |
| CNBG_1745 | Cda1 | 0.0 | 4.54E-37 | 4.58E-36 |
| CNBG_0964 | Cda2 | 2.07E-35 | 0.0 | 3.63E-29 |
| CNBG_0806 | Cda3 | 1.01E-37 | 2.34E-34 | 0.0 |

### *C. gattii* R265 cells produce significantly higher amount of chitosan in the cell wall compared to *C. neoformans* under YPD growth conditions

We have shown for *C. neoformans* that the amount of cell wall chitosan significantly influences host immune response during infection (26, 29). Various studies have demonstrated that R265 elicits different types of immune response either in the host or when incubated with immune cells under *in vitro* conditions when compared to the response induced by *C. neoformans* cells (22–25). Therefore, we were curious to see if there is a difference in the amount of chitosan between KN99, a hypervirulent strain of *C. neoformans* (31) and R265. To determine the amount of chitosan, we grew both KN99 and R265 cells in YPD at 30°C. We have previously reported the specific affinity of an anionic dye Eosin Y to chitosan in *C. neoformans* (28). Therefore, we stained wild-type *C. neoformans* KN99 and *C. gattii* R265 cells after five days of culture. As shown in Fig. 1A, we observed a dramatic increase in the binding of Eosin Y to R265 cells compared to KN99 cells. This difference was further quantified by measuring the mean fluorescence intensity (MFI) per cell using ImageJ (Fig. 1B). The MFI/cell of KN99 was 32.9 compared to 62.9 for R265. We then measured the total amount of chitin and chitosan biochemically employing the 3-methyl-2-benzothiazolinone hydrazone (MBTH) assay as described in Materials and Methods. Wild-type KN99 and R265 cells were grown for 1-5 days. At different days of growth, the cellular chitosan was quantified by the MBTH assay. As shown in Figs. 1C and 1D, at day two, R265 cells started to show increased amount of chitosan compared to KN99 and this increase peaked at day three of growth before it started to decrease. At its peak, the amount of chitosan as expressed as nmoles of glucosamine per mg dry weight of the cell wall material in the R265 cells had increased by three fold compared to the cells of KN99. The levels of chitin in the R265 cells remained almost constant throughout the growth period and their levels were comparable to KN99 cells. The sharp increase in the amount of chitosan observed in R265 on day three was not observed in KN99 cells which showed a slight increase in the chitosan amount (Fig.1C and 1D). These data indicate that outbreak R265 cells produce significantly higher amount of chitosan compared to those produced by KN99 cells under YPD growth conditions.

**Figure 1.**
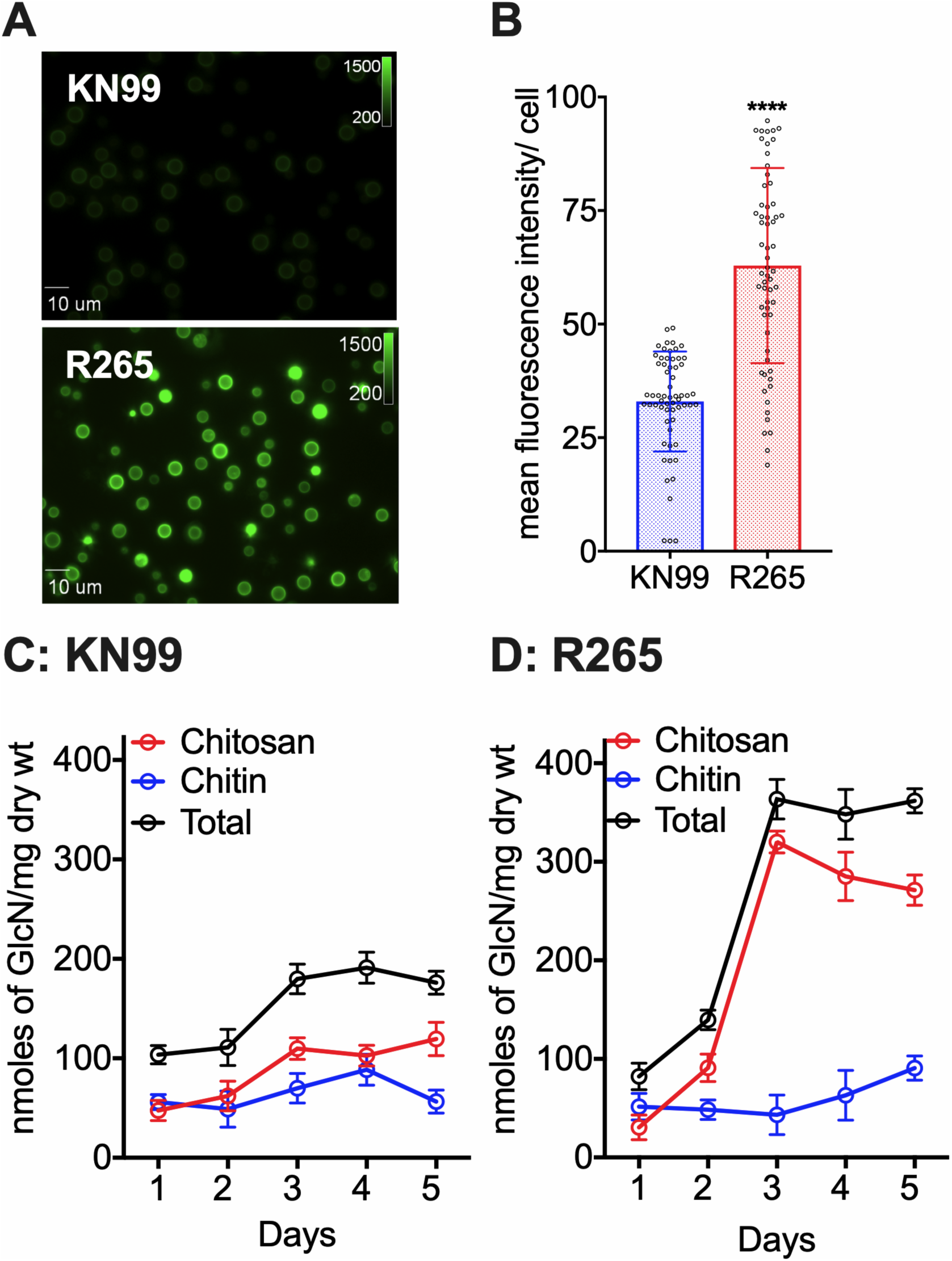
*C. gattii* R265 cells produce significantly higher amounts of chitosan in the cell wall compared to *C. neoformans* under YPD growth conditions. (**A**) Wild-type strains, R265 and KN99 were grown in YPD for five days at 30°C and stained with Eosin Y to detect cell wall chitosan. Staining intensity was assessed using epifluorescence microscopy with identical exposures for all images. (**B**). Fluorescent levels for 60 individual cells (represented in panel A) were quantified using ImageJ (Fiji). The two-tailed unpaired t test with Welch’s correction was used to compare means values of the wild-type. Means represent the fluorescent (Fluor.) intensity levels from three independent experiments (n = 3). ****, P < 0.0001. Error bars represent standard errors of the means. (**C**) Quantitative determination of cell wall chitosan and chitin of KN99 by the MBTH assay. Cells were grown in YPD for one to five days, collected, washed and used for the assay. Data represents the average of three biological experiments and are expressed as ηmoles of glucosamine per mg dry weight of yeast cells. (**D**) Quantitative determination of cell wall chitosan and chitin of R265 were determined same as panel (**C**).

### *C. gattii* R265 cells produce significantly higher amount of chitosan compared to *C. neoformans* under host conditions

To determine if the increased amount of chitosan was also seen under host conditions, we first measured chitosan of KN99 and R265 cells cultured in tissue culture conditions and then for Cryptococcus isolated from infected mouse lungs. Wild-type KN99 and R265 cells were first grown in YPD medium at 30°C and then transferred to RPMI 1640 medium containing 10% fetal bovine serum (FBS), 5% CO^2^ at 37°C for 6 days. The cells were then harvested and the MBTH assay was used to quantify the chitosan content. As shown in Fig. 2A, R265 cells have a mean value of 63.9 versus 30.2 of KN99 cells as expressed as nmoles of glucosamine sugar per mg dry weight of the cell wall material. Next, chitosan was determined after isolating the yeast cells from infected mouse lungs. Similar to YPD and RPMI culture conditions, R265 cells showed significantly higher cell wall chitosan compared to KN99 cells with mean values of 213.9 and 78.3, respectively, as expressed as nmoles of glucosamine sugar per 10^8^ cells (Fig. 2B). These results suggest that the increase in cell wall chitosan of *C. gattii* R265 over *C. neoformans* KN99 is a phenotype that is preserved when these strains are grown in culture and during mouse infection.

**Figure 2.**
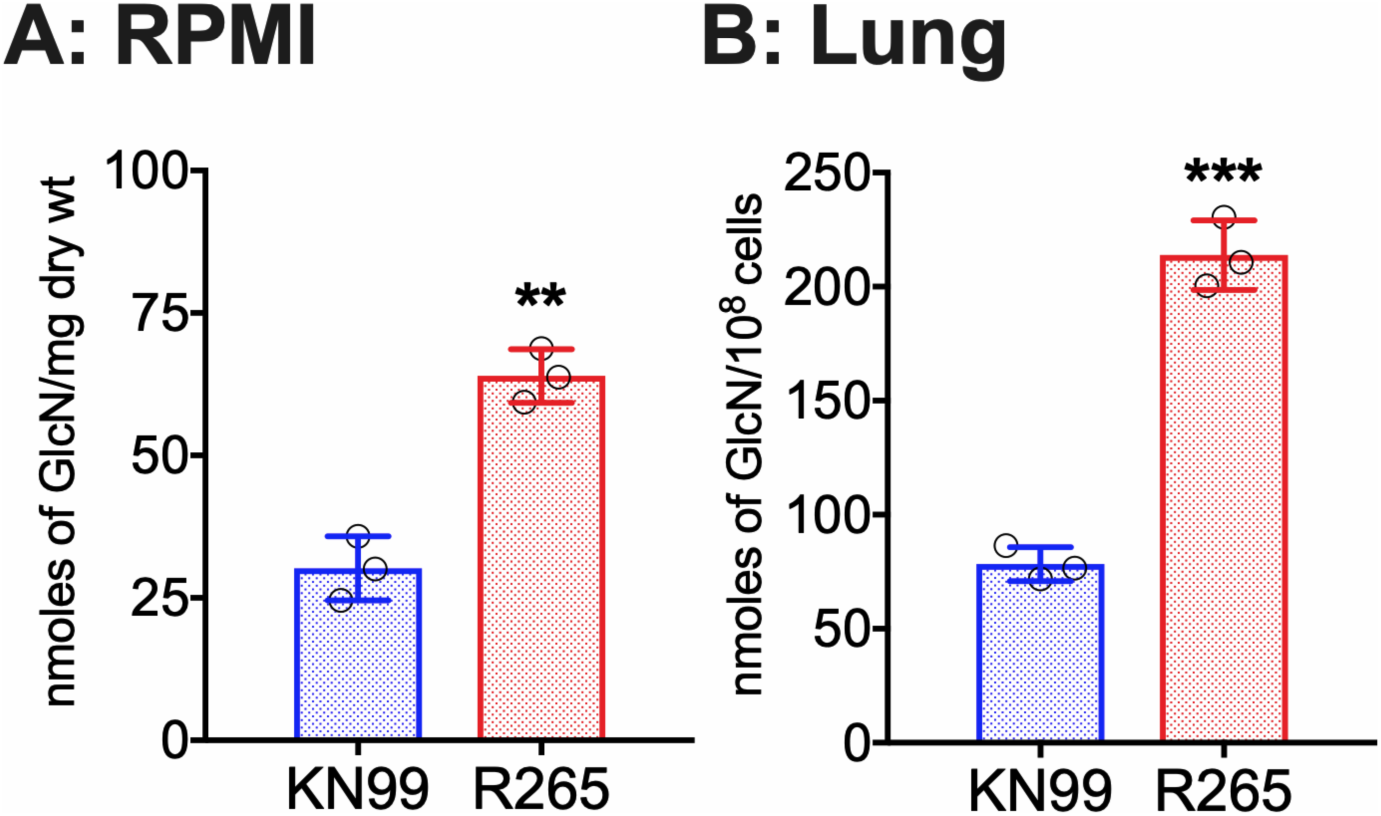
*C. gattii* R265 cells produce significantly higher amount of chitosan in the cell wall compared to *C. neoformans* under host conditions. (**A**) Chitosan levels of strains grown in RPMI containing 10% FBS and 5% CO^2^ at 37°C for five days. Strains were grown in YPD for two days. Yeast cells were harvested, washed with PBS and inoculated at 500 cells/µL in RPMI containing 10% FBS and incubated for five days at 37°C in the presence of 5% CO^2^. At the end of incubation, chitosan was measured by the MBTH assay and expressed as ηmoles of glucosamine per mg dry wt cells. Data represent the average of three biological experiments. (**B**) Chitosan levels of strains growing in the murine lung. Mice (CBA/J; three per group) were intranasally inoculated with 10^7^ CFU of each strain. On day seven PI, lungs were excised, homogenized and the lung tissue was removed by alkaline extraction leaving the fungal cells to be harvested, counted and subjected to the MBTH assay. Data are expressed as ηmoles of glucosamine per 10^8^ cells. Significant differences between the groups were compared by two-tailed unpaired t test with Welch’s correction. Error bars represent standard errors of the means. *** P < 0.0062 and ** P < 0.001.

### The deletion of R265, Cda1 displays a decrease in cell wall chitosan under YPD growth conditions

We generated single CDA deletion mutants of R265 by biolistic transformation. The strains generated and used in this study are listed in Table 2. The deletion cassettes for each gene deletion were generated by overlap PCR and were biolistically transformed into R265 cells. The primers used to generate these deletion cassettes are listed in supplementary Table 1. All the isolates were characterized by diagnostic PCR screening and southern blot hybridization. After growing the strains in YPD for five days, we measured chitosan by MBTH assay. We found that deletion of the *CDA1* gene decreased the chitosan amount by 33% compared to wild-type R265. However, deletion of either *CDA2* or *CDA3* did not affect the total amount of chitosan, as shown in Fig.3. This is different from what we have previously described for *C. neoformans*, where there was no significant difference in chitosan in any of the single CDA deletion strains (28, 30). This suggests that there are differences in the regulation of chitin deacetylation between the two species, with R265 Cda1 having a more significant role in chitosan production when the cells were cultured in YPD.

**Figure 3.**
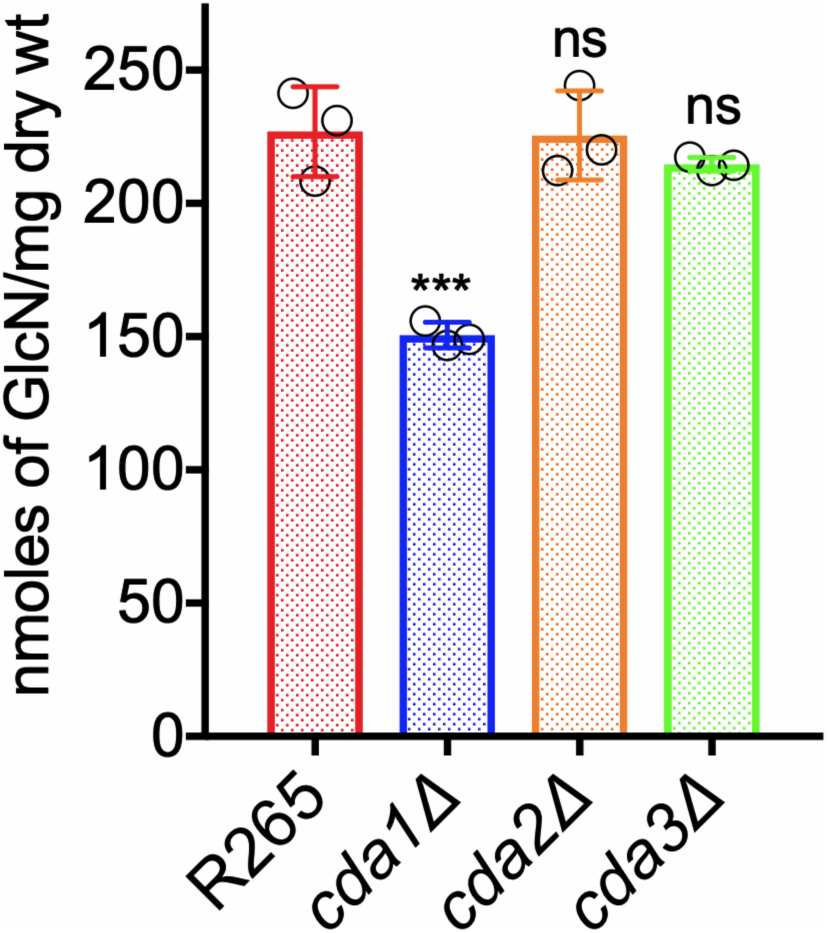
The deletion of R265, *CDA1* displays a decrease in cell wall chitosan under YPD grown conditions. Chitosan levels of strains grown in YPD. Strains were grown in YPD for five days. The amount of chitosan in the cell wall of the strains was quantified by the MBTH assay. Data represents the average of three biological experiments with two technical replicates and are expressed as ηmoles of glucosamine per mg dry weight of yeast cells. Significant differences between the groups were compared by one-way ANOVA followed by Dunnett’s multiple comparison test. *** p <0.0002 comparing wild-type R265 with any other strain.

**Table 2.**
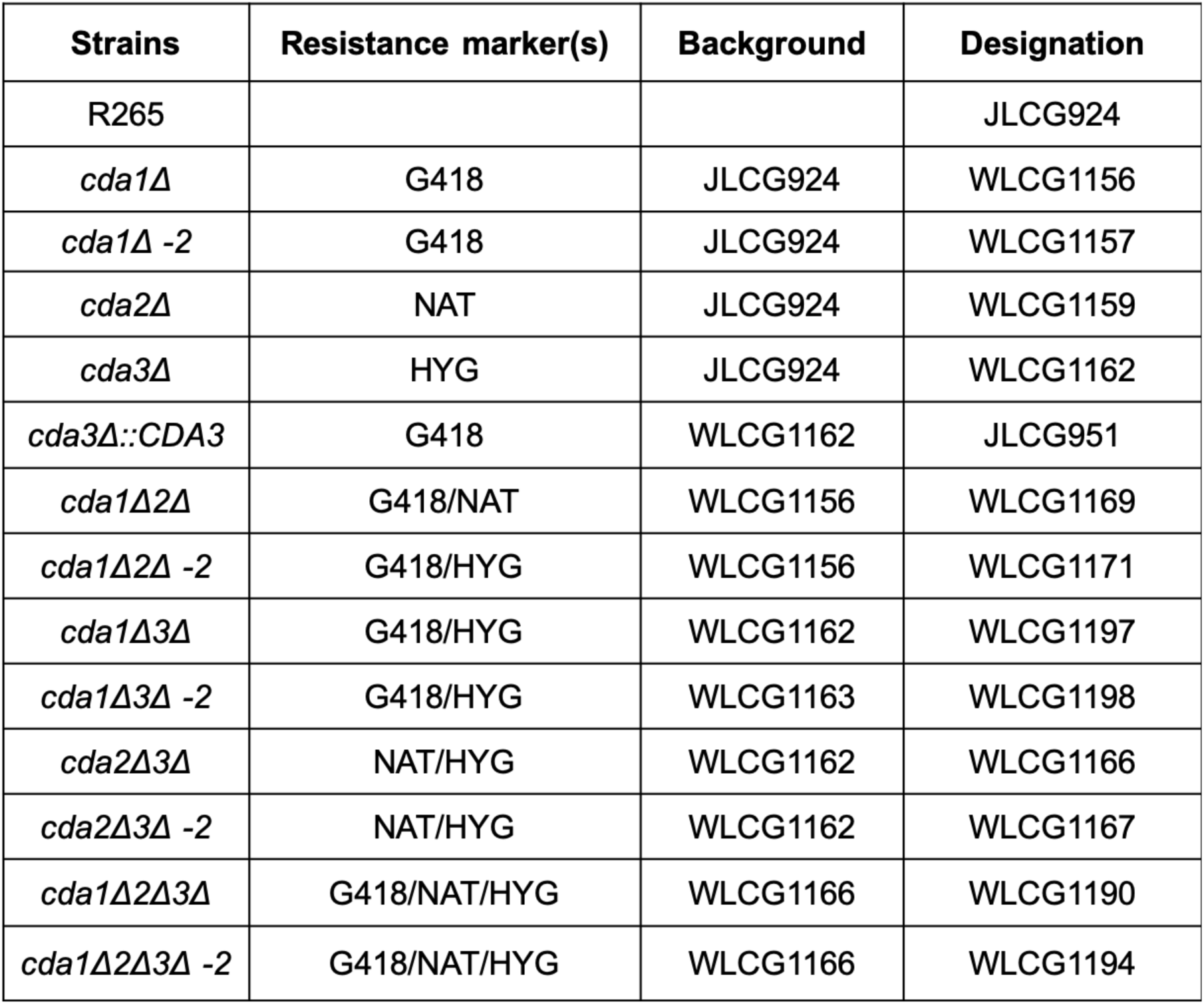
Strains used and generated in this study.

### R265 Cda3 is critical for fungal virulence

Next, we assessed the CDA single-deletion mutant strains for virulence by employing the murine intranasal infection model. We infected CBA/J mice with 10^5^ wild-type cells or with cells of the corresponding single CDA deletion strains. The virulence was assayed as described in the Materials and Methods. We found that deletion of either *CDA1* or *CDA2* did not affect the virulence (Fig. 4A). The virulence of *cda1Δ* was reproduced using a second isolate and is shown in Supplementary Fig. 2 to further confirm the absence of a role for Cda1 in the virulence of *C. gattii* R265. Interestingly, we found that R265 Cda3 is essential for fungal virulence. The virulence defect of the *cda3Δ* was completely restored in a *CDA3* complemented strain (Fig. 4A). Since different mouse models show varying degrees of sensitivity to *Cryptococcus* infection, we wanted to verify whether the associations between the absence of CDA genes, amount of cell wall chitosan and the virulence of the specific CDA deletion strain can be recapitulated in C57BL/6 mice, which is routinely employed for diverse immunological studies using readily available mutants. Similar to the results obtained with CBA/J mice, we found that only R265 *cda3Δ* is specifically avirulent when tested following orotracheal inoculation of 10e4 CFU of yeast (Fig. 4B). The avirulent phenotype in both mouse strains was accompanied by the complete clearance of the mutant strain from the infected host at the end point of the survival study (Supplementary Fig. 3). This was rather surprising because we have previously seen that for KN99, Cda1 plays a major role in fungal virulence without the contribution of Cda3 to pathogenesis and that *C. neoformans cda1Δ* mutant cells persist in the mouse lungs at low levels, even though the strain doesn’t cause disease (30).

**Figure 4.**
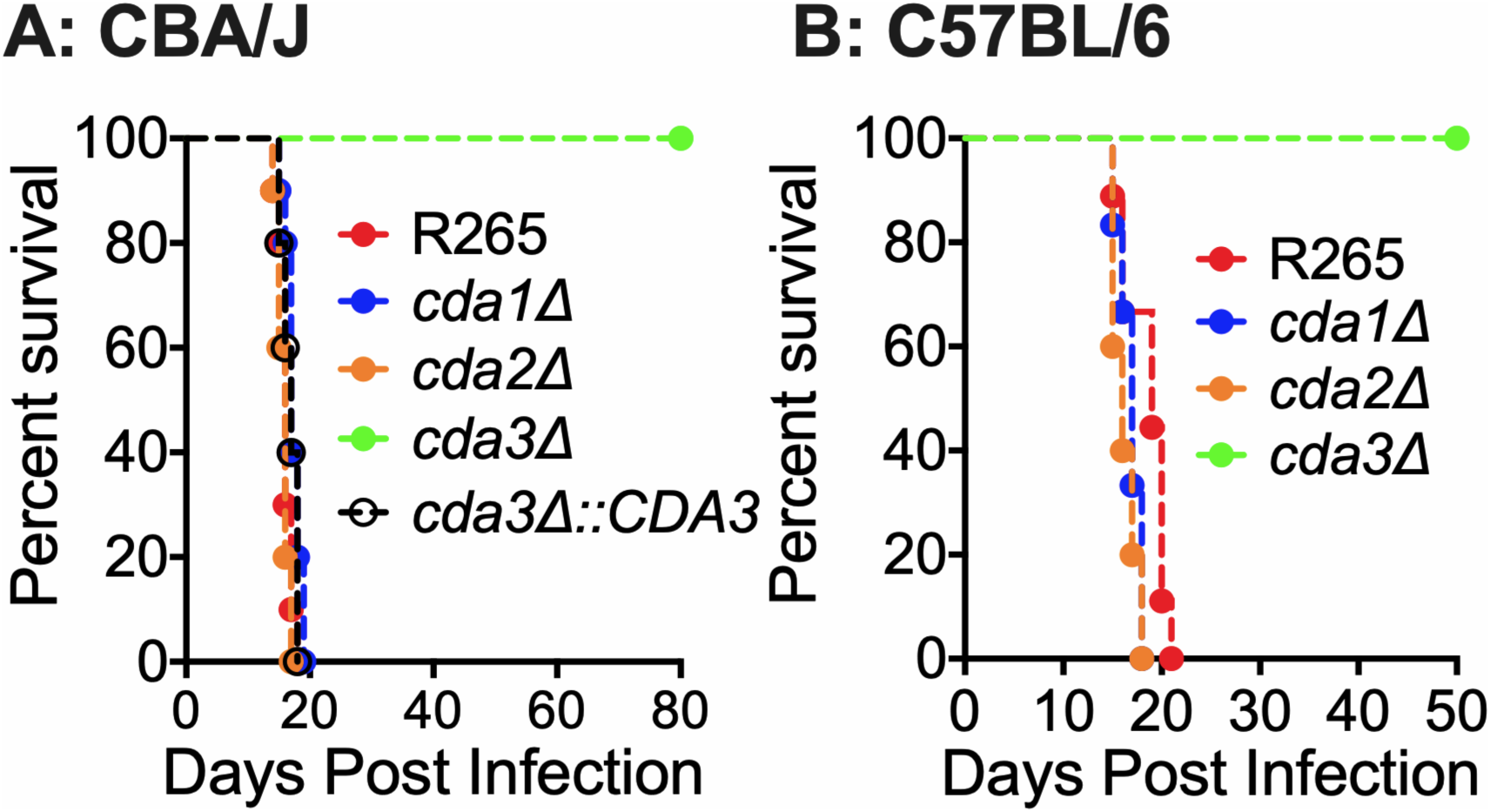
*C. gattii cda3Δ* displays severely attenuated virulence in CBA/J and C57BL/6 mouse model of infection. (**A**) CBA/J mice (4-6 weeks old, female) were infected intranasally with 10^5^ CFU of each strain. Survival of the animals was recorded as mortality of mice for 80 days PI. Mice that lost 20% of the body weight at the time of inoculation were considered ill and sacrificed. Data is representative of two independent experiments with five animals for each strain. (**B**) C57BL/6 mice (4-6 weeks old, female) were infected with 10^4^ CFU of each strain by intratracheal inoculation. Survival of the animals was recorded as mortality of mice for 50 days PI. Mice that lost 20% of the body weight at the time of inoculation were considered ill and sacrificed. Data is representative of one experiment with 10 animals for each strain. Virulence was determined using Mantel-Cox curve comparison with statistical significance determined by log-rank test. p<0.0001 comparing KN99 with *cda3Δ*.

The virulence phenotype of the single CDA deletion strains followed their ability to grow in the infected lung. As shown in Fig. 5, we found that CBA/J mice infected with either *cda1*Δ or *cda2*Δ strains had lung fugal burden similar to that of R265 on day 14 and 21 PI. On the other hand, mice infected with the *cda3*Δ strain showed slow and gradual clearance of the mutant strain. The fungal growth in the lungs of the mice infected with the *cda3*Δ strain was restored to wild-type R265 levels in a *CDA3* complemented strain (Fig. 5; *cda3*Δ::*CDA3*).

**Figure 5.**
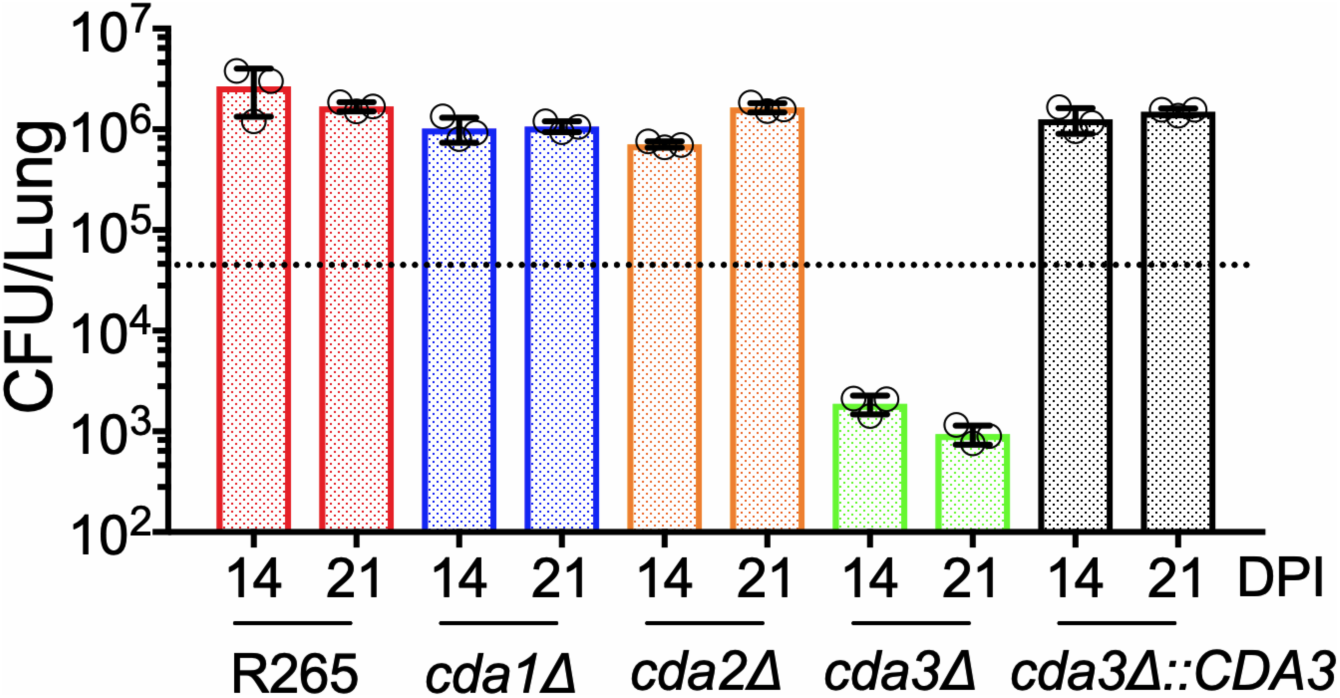
There is a slow but gradual clearance of *C. gattii cda3Δ* in CBA/J mice. Fungal burden in the lungs of CBA/J mice infected with indicated strains at different days after infection. Data are from three mice per group at each time point. The dashed line indicates the CFU of the initial inoculum for each strain.

### Fungal virulence of different CDA deletion strains is directly correlated with their ability to produce chitosan under host-mimicking growth conditions

Recently, we have shown that in *C. neoformans*, Cda1 is essential for fungal virulence, and this avirulent phenotype is associated with the inability of the mutant to produce wild-type levels of chitosan when grown under host-mimicking conditions, such as RPMI 1640 medium containing 10% FBS, 5% CO^2^, and 37°C (30). Therefore, we wanted to test whether such defects are also responsible for the avirulent phenotype of R265 mutants that do not harbor the *CDA3* gene in the genome. As shown in Fig. 6, the loss of Cda3 resulted in a 77% decrease in the amount of chitosan produced compared to wild-type R265. Even though the amount of chitosan in the *cda1Δ* strain showed 33% reduction compared to wild-type R265 (Fig. 6), this decrease was similar to what we observed when the cells were grown in YPD culture conditions as well (Fig.3). These data indicate that in R265, Cda1 mediated deacetylation of chitin is not influenced by the growth conditions and is not critical for virulence. However, Cda3 is responsible for the majority of the deacetylation either in RPMI culture conditions or in infected mice and thus contributes significantly to fungal proliferation in the host. Taken together, these results confirmed the importance of cell wall chitosan to fungal virulence and suggests that a certain threshold amount of chitosan needs to be maintained in the cell wall to sustain infection.

**Figure 6.**
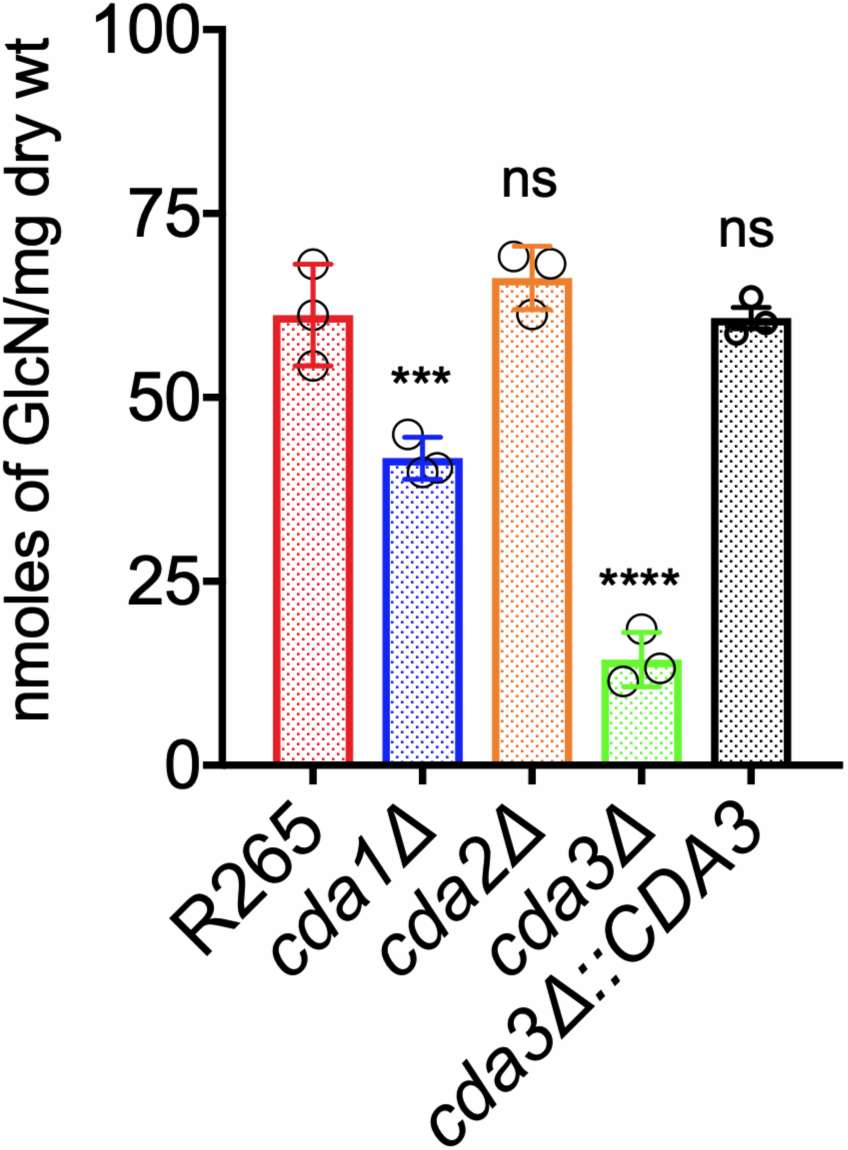
*C. gattii* Cda3 plays a major role in the synthesis of chitosan under host infection conditions. Chitosan levels of strains grown in RPMI containing 10% FBS and 5% CO^2^ at 37°C for five days. The indicated strains were grown in YPD for 48 hours. Yeast cells were harvested, washed with PBS, and inoculated at 500 cells/μL in RPMI containing 10% FBS and incubated for five days at 37°C in the presence of 5% CO^2^. At the end of incubation, chitosan was measured by the MBTH assay and expressed as ηmoles of glucosamine per milligram (dry weight) of cells. Data represent the averages for three biological experiments. Significant differences between the groups were compared by one-way ANOVA, followed by Dunnett’s multiple-comparison test (*** P < 0.0002, * P < 0.0280 comparing KN99 with any other strain; ns, not significant).

### Double and triple CDA mutants of R265 displayed varied amounts of chitosan under YPD and host-mimicking conditions

We generated three double CDA deletion strains and a triple CDA gene deletion strain in R265 by biolistic transformations as indicated in Table 2. We subjected these strains to chitosan quantification after growing them either in YPD or in host-mimicking conditions of RPMI containing 10% FBS in the presence of 5% CO_2_ at 37°C. When we compared the chitosan amounts among the CDA double deletion strains, we found that the cell wall chitosan amount was significantly decreased in all the double deletion strains grown under YPD culture condition (*cda1Δ2Δ*, *cda1Δ3Δ* or *cda2Δ3Δ* in Fig. 7A). The decrease in the chitosan amount was more pronounced in the absence of Cda1 in combination with either Cda2 or Cda3: 63% reduction for the *cda1Δ2Δ* strain and 54% reduction for the *cda1Δ3Δ* compared to wild-type R265. The amount of chitosan in *cda2Δ3Δ* showed an only 26% reduction compared to wild-type R265.

**Figure 7.**
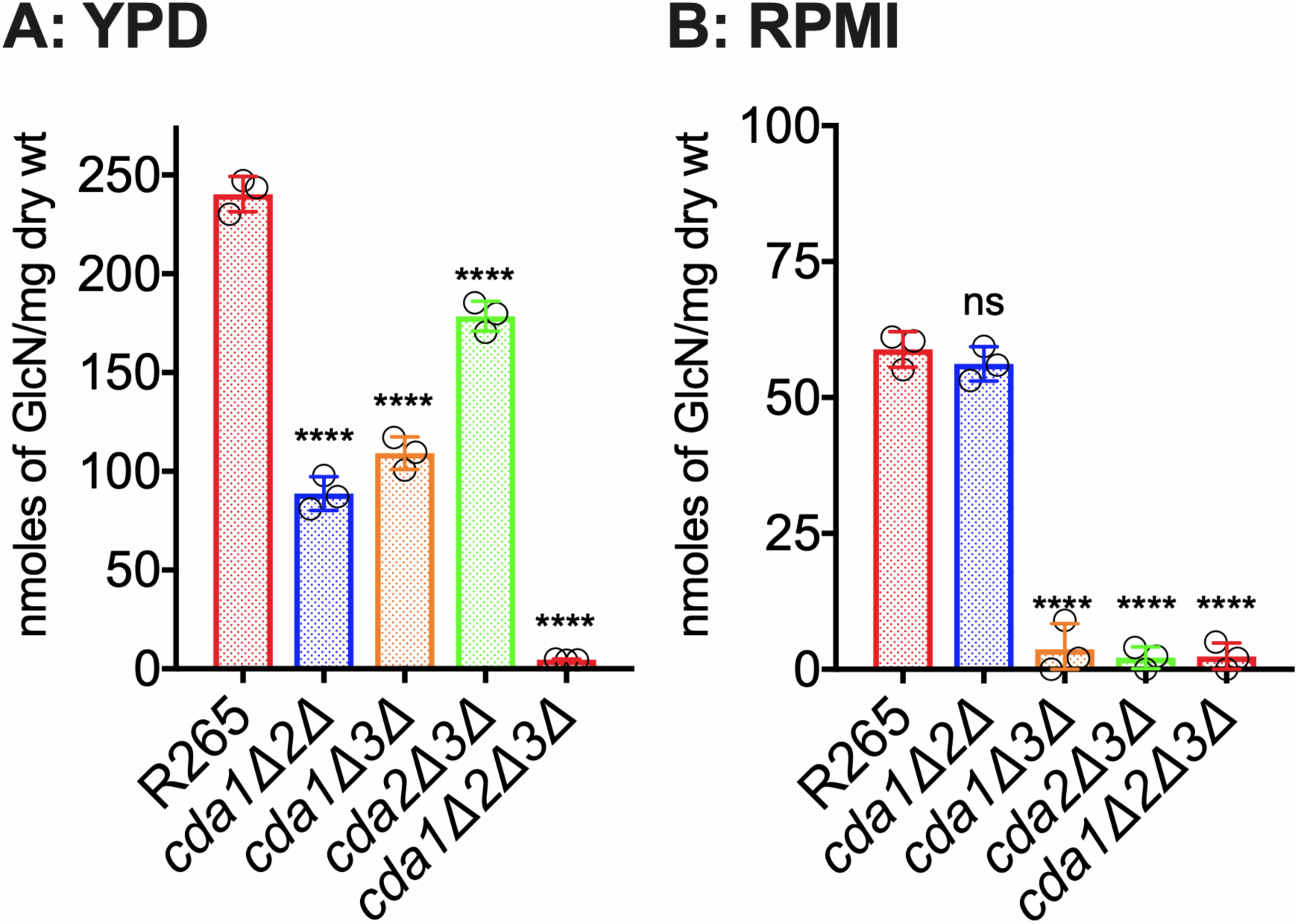
The deletion of *C. gattii CDA1*, *CDA2*, and *CDA3* results in a strain that is completely chitosan-deficient. *C. gattii* Cda1 in combination with Cda2 or Cda3 plays a major role in chitosan synthesis in vegetative growing conditions. While *C. gattii* Cda3 in combination with Cda1 or Cda2 results in a stain that is completely chitosan-deficient. (**A**) Chitosan levels of strains grown in YPD. The indicated strains were grown in YPD for five days. The amount of chitosan in the cell wall of the strains was quantified by the MBTH assay. Data are the averages for three biological experiments and are expressed as ηmoles of glucosamine per milligram (dry weight) of yeast cells. (**B**) Chitosan levels of strains grown in RPMI containing 10% FBS and 5% CO2 at 37°C for five days. The indicated strains were grown in YPD for 48 hours. Yeast cells were harvested, washed with PBS, and inoculated at 500 cells/μL in RPMI containing 10% FBS and incubated for 5 days at 37°C in the presence of 5% CO^2^. At the end of incubation, chitosan was measured by the MBTH assay and expressed as ηmoles of glucosamine per milligram (dry weight) of cells. Data represent the averages for three biological experiments. Significant differences between the groups were compared by one-way ANOVA, followed by Dunnett’s multiple-comparison test (**** P < 0.0001, comparing KN99 with any other strain; ns, not significant).

These results are consistent with the major role of Cda1 in chitosan production in R265 when the cells were grown under YPD conditions (Fig. 3). When we deleted all the three CDAs to generate a *cda1Δ2Δ3Δ* strain, the chitosan amount decreased almost to negligible amounts, suggesting that in spite of differences in the role of individual CDAs between KN99 and R265, the deletion of all three CDAs was sufficient to render the mutant chitosan-deficient (Fig. 7A). When the strains were grown under host-mimicking conditions (RPMI+10% FBS grown with CO_2_ at 37°C), any strain in which *CDA3* was deleted in combination with either *CDA1* or *CDA2* (*cda1Δ3Δ* or *cda2Δ3Δ* in Fig. 7B) produced negligible amounts of chitosan similar to the *cda1Δ2Δ3Δ* strain, further indicating the importance of R265 Cda3 in the production of chitosan in the host and its subsequent effect on fungal virulence.

### Chitosan-deficient R265 cells were sensitive to cell wall stressors and had normal capsule and melanin producing abilities

We subjected the single CDA deletion strains and the triple CDA deletion strain of R265 to a panel of cell wall stressors (0.005% SDS, 1 mg/ml Calcofluor white, 0.5 mg/ml caffeine, 0.4% Congo Red) added to YPD agar medium, which are routinely employed to determine cell wall integrity. As shown in Supplementary Fig. 3, only the chitosan-deficient *cda1Δ2Δ3Δ* strain was sensitive to the various cell wall stressors. None of the CDA deletion mutants were sensitive to temperature when their growth at 30°C was compared to the growth at 37°C on YPD agar medium. The deletion of CDA genes in R265 either individually or in combination did not affect their ability to either produce capsule (Supplementary Fig. 4) or melanin as shown in Supplementary Fig. 5. In contrast to *C. neoformans* mutants (26), none of the CDA mutants of R265 displayed “leaky melanin” phenotype.

### R265 *CDA1* and *CDA2* are dispensable for virulence even when both genes were deleted

We next subjected the double and triple CDA gene deletion strains to tests of fungal virulence either by intranasal infection of CBA/J or by orotracheal inoculation of C57BL/6. We found that deletion of *CDA1* and *CDA2* did not affect the virulence in either CBA/J or C57BL/6 (*cda1Δ2Δ*; Fig. 8A-B). Consistent with the role of R265 Cda3 in virulence, double CDA deletion strains in which *CDA3* is deleted in combination with either *CDA1* or *CDA2* showed a major defect in virulence (*cda1Δ3Δ* and *cda2Δ3Δ* Fig. 8A-B). These results further point to the importance of just Cda3 in fungal virulence for R265. The mutant strain devoid of all the three CDA genes (*cda1Δ2Δ3Δ*) was completely avirulent (Fig. 8A-B). For all the mutants, the avirulent phenotype of the mutant strains was accompanied by the inability of the different CDA mutant strains to proliferate or maintain in the infected murine lung as revealed by their gradual clearance (data not shown). To further ascertain these results, we subjected second independent isolate of each mutant to fungal virulence studies and obtained nearly identical results (Supplementary Fig. 6).

**Figure 8.**
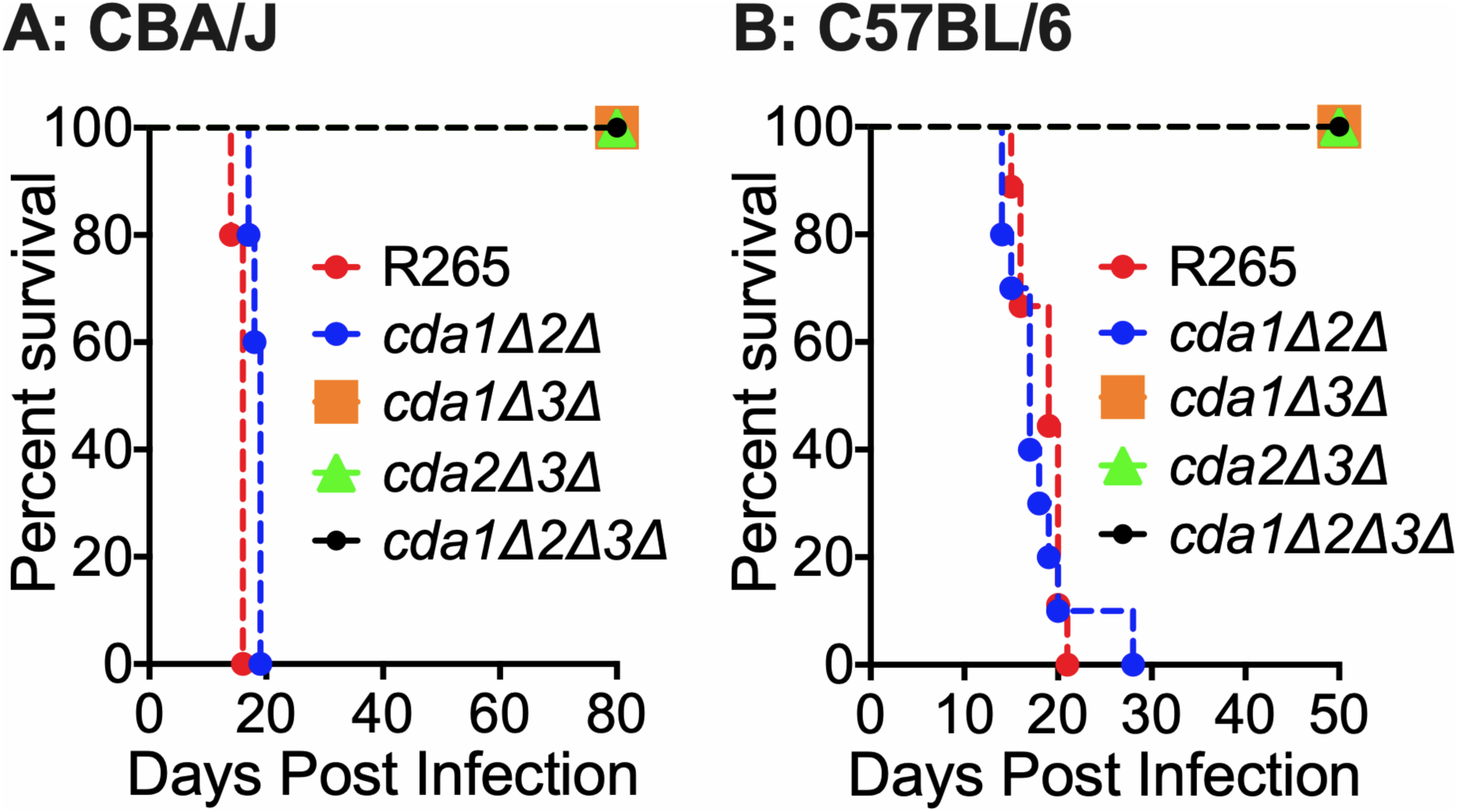
Deletion of *C. gattii CDA3* in combination with any of the other two CDAs results in severe attenuation of virulence in CBA/J and C57BL/6 mouse models of infection. (**A**) CBA/J mice (6-8 weeks old, female) were infected intranasally with 10^5^ CFU of each strain. Survival of the animals was recorded as mortality of mice for 80 days PI. Mice that lost 20% of the body weight at the time of inoculation were considered ill and sacrificed. Data is representative of two independent experiments with five animals for each strain. **B**) C57BL/6 mice (4-6 weeks old, female) were infected intratracheally with 10^4^ CFU of each strain. Survival of the animals was recorded as mortality of mice for 50 days PI. Mice that lost 20% of the body weight at the time of inoculation were considered ill and sacrificed.

### Vaccination with the chitosan-deficient R265 cells confer partial protection to subsequent challenge with the virulent wild-type R265

For *C. neoformans*, we observed that mice infected with 10^7^ CFU of chitosan-deficient *cda1Δ2Δ3Δ* strain resulted in the clearance of the mutant. That clearance was accompanied by the induction of a robust protective response to a subsequent infection with wild-type, fully virulent KN99 (29). This protective response was only observed when mice were vaccinated with 10^7^ CFU of *cda1Δ2Δ3Δ,* while vaccination with either 10^6^ or 10^5^ CFU did not generate a protective response (29). Therefore, since the complete clearance of the chitosan-deficient *cda1Δ2Δ3Δ* strain is also observed for mutants generated in the R265 background, we wanted to determine whether this clearance of the mutant strain induces protective immunity to R265 infection. For this, we vaccinated naïve mice (CBA/J) with either 10^5^, 10^6^ or 10^7^ CFU of live preparations of *cda1Δ2Δ3Δ* by intranasal inhalation. After 40 days PI, we challenged them with 10^5^ CFU of R265. Two independent isolates of *cda1Δ2Δ3Δ* (*cda1Δ2Δ3Δ*−1 and *cda1Δ2Δ3Δ*−2) were used for this study. As shown in Fig. 9A, vaccination with either of the chitosan-deficient *cda1Δ2Δ3Δ* isolate conferred only partial protection to subsequent infection with R265. The control mice that received PBS alone succumbed to R265 infection with a median survival time of 20 days PI. However, the mice vaccinated with live-preparation of *cda1Δ2Δ3Δ* had a median survival of 30-40 days PI upon secondary challenge infection with R265. This protective immunity required a minimal dose of 10^7^ CFU of *cda1Δ2Δ3Δ* for vaccination (Fig. 9A), as observed for *cda1Δ2Δ3Δ* of KN99 (29). Next, we wanted to determine whether heat-killed (HK) preparation of *cda1Δ2Δ3Δ* induces protective immunity. We confirmed that incubating live R265 (either wild-type or the corresponding *cda1Δ2Δ3Δ* mutant) at 70°C for 15 min was sufficient to kill both strains by plating them for CFUs onto YPD agar. Then we vaccinated mice intranasally with 10^7^ CFU of HK preparation of either R265 or *cda1Δ2Δ3Δ* waited 40 days and challenged them with 10^5^ CFU of R265. The mice vaccinated with either PBS or with HK wild-type R265 succumbed to R265 infection with a median survival time of 20 days PI. However, the mice vaccinated with the HK *cda1Δ2Δ3Δ* had a median survival of 33 days PI (Fig. 9B) similar to that observed for vaccination with the live *cda1Δ2Δ3Δ* (Fig. 9A). When the vaccination and protection experiment was done in C57BL/6 mice using HK *cda1Δ2Δ3Δ* as a vaccine, mice had a median survival of 41.5 days compared to 26 days of median survival for unvaccinated mice (Supplementary Fig.7).

**Figure 9.**
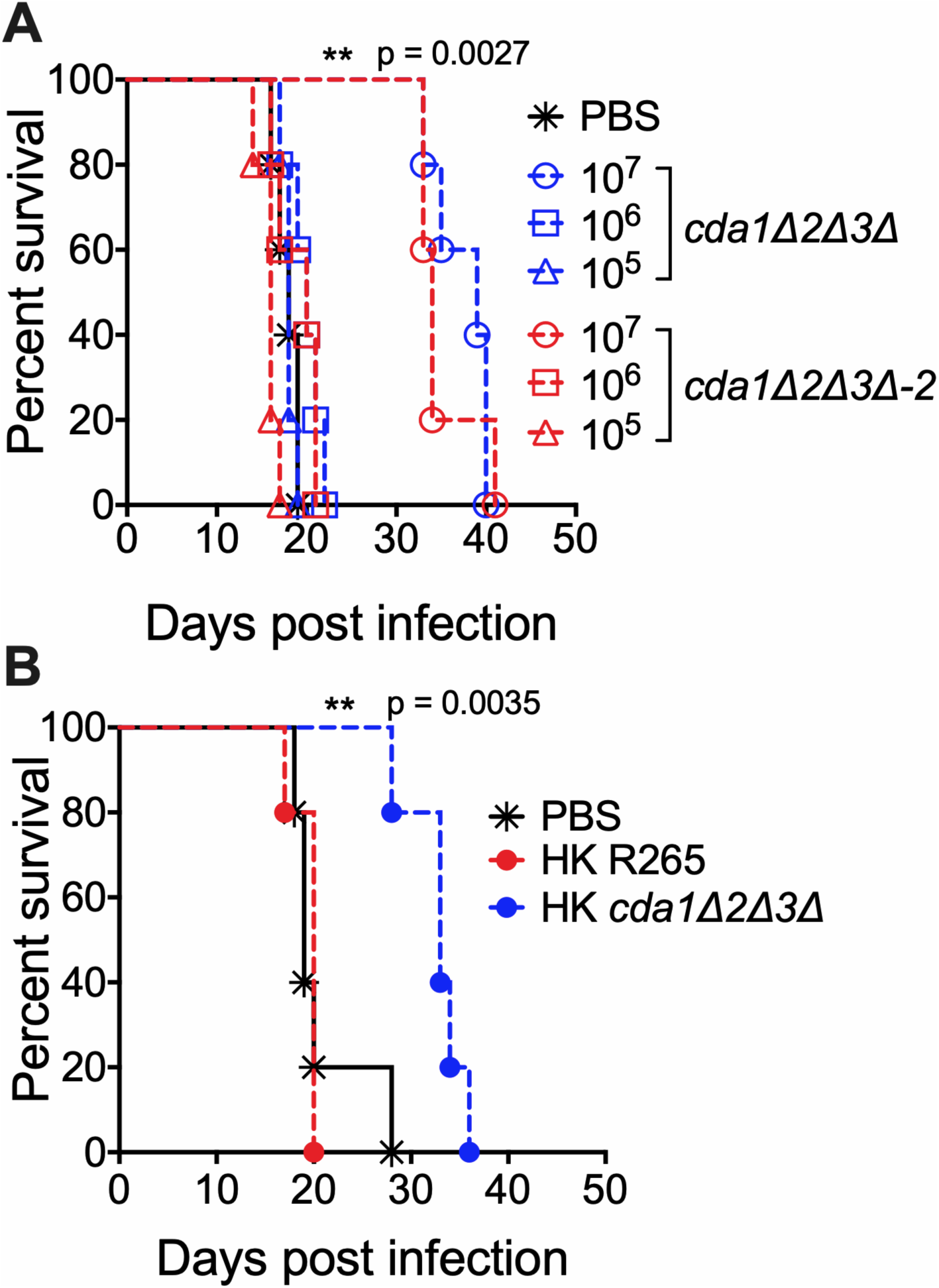
Vaccination of CBA/J mice with 10^7^ CFU of live or heat-killed (HK) *cda1Δ2Δ3Δ* cells conferred attenuated protective immunity to subsequent infection with wild-type R265 *C. gattii* cells. (**A**) Mice were immunized with either 10^7^, 10^6^, or 10^5^ live CFU of *cda1Δ2Δ3Δ* through inhalation. PBS-inoculated mice served as control. Animals were left for 40 days to resolve the infection. Subsequently, both groups of mice were challenged with 50,000 CFU of virulent R265 cells. Virulence was recorded as mortality of mice. Mice that lost 20% of starting body weight at the time of inoculation were considered to be moribund and were sacrificed. The percentage of mice that survived was plotted against the day p.i. Each survival curve is the average of two independent experiments that had five mice per experimental group. (**B**) Mice were immunized with an inoculum of HK cells with a dose equivalent to 10^7^ CFU of either the wild-type R265 or the *cda1Δ2Δ3Δ* strain. Control mice were inoculated with PBS. After 40 days, mice were challenged with 50,000 CFU of wild-type R265 cells. Survival of the animals was recorded as described above. The data shown are the average results from two experiments with five mice per experimental group.

## Discussion

Chitosan is one of the principle components of the *C. neoformans* cell wall. In its absence, yeast cells display a severe budding defect resulting in irregular shaped, often clumped cells with significant sensitivity to various cell wall perturbing agents (28). More importantly, absence of chitosan makes them completely avirulent in a mammalian host as the chitosan-deficient cells stimulate robust host immune responses, which in turn leads to rapid clearance of infection (26, 29). The outbreak strain of *C. gattii*, R265 is a hyper-virulent strain that has been reported to possess several key features that enabled it to cause symptomatic infection, even in immune competent individuals (10, 32, 33). Its higher rate of intracellular proliferation with increased resistance to oxidative stress, its ability to dampen host immune response and its potential to inhibit the maturation of specific immune cells all may contribute to its virulence (19, 22–25, 34, 35). A higher amount of chitosan in R265 than *C. neoformans* KN99 may efficiently shield surface exposed PAMPs from being recognized by the host immune system, thereby limiting the intensity and the complexity of the host immune response. *C. neoformans* and *C. gattii* share diverse ecological niches. Moreover, *C. gattii R265* is predominantly detected in the environment and strictly associated with plants where they encounter chitinases from soil microbes and from plant hosts respectively (32). Extensive deacetylation of chitin to form chitosan may provide greater resistance to environmental chitinases thereby increasing its fitness in the environment.

*C. neoformans* Cda1 plays an important role in the deacetylation of chitin especially during host infection (30). Among the individual *C. neoformans* CDA deletion strains, we did not observe phenotypic differences when cultured in YPD medium suggesting the redundancy in their function (28). However, deletion of Cda1 alone caused a dramatic avirulent phenotype in virulence studies using CBA/J mice (30). Unlike *C. neoformans*, Cda1 of *C. gattii* seems to play an important role in the deacetylation when grown in YPD medium since its deletion caused a 34% reduction in the amount of cell wall chitosan (Fig. 3). The deletion of either *CDA2* or *CDA3* did not significantly reduce the chitosan amount when grown in YPD. However, when grown under host mimicking conditions of RPMI with 10% FBS, 5% CO_2_ and at 37°C, *C. gattii* Cda3 played an important role in the deacetylation of chitin, since its deletion in *cda3*Δ caused a 76% reduction in the amount of chitosan. On the other hand, the decrease in the amount of chitosan in the *cda1*Δ when grown in RPMI conditions was around 32%, which is similar to what we observed for YPD culture conditions suggesting a unique role of R265 Cda3 in deacetylating chitin in host mimicking conditions. For our initial animal virulence experiments, we chose two independent isolates of *cda1*Δ expecting that they will mimic the avirulent phenotype observed in *C. neoformans* (30). However, we were surprised to see that for *C. gattii* R265, deacetylation of chitin during host infection was even more dependent on Cda3 than Cda1. The avirulent phonotype of *cda3*Δ was further confirmed by the *CDA3* complemented strain in CBA/J mouse. In spite of its role in chitosan production in YPD and host mimicking media, the decreased levels of chitosan in *cda1*Δ did not affect its virulence (Fig. 4). It may be that the levels of chitosan present in the *cda1*Δ during infection is still sufficient enough for promoting pathogenesis or the pattern of deacetylation in the chitosan produced by Cda1 may be different from that of Cda3. This differences in the molecular structure of chitosan may have contributed to the observed differences in the virulence potential between different CDA deletion mutant strains.

All three CDAs contribute to chitosan production in R265 while growing in YPD since deletion of two *CDA* genes in combination significantly reduced cell wall chitosan (Fig 7). This again was reflected in the virulence phenotype of the double and triple CDA deletion strains. When *CDA3* was deleted in combination with any one or both of the other CDA genes the resulting strains had substantially reduced chitosan when grown in host-mimicking conditions and were avirulent. Any strain in which *CDA3* is deleted was avirulent in both CBA/J and C57BL/6 mouse strains and was cleared from the infected lung. Deletion of all three CDAs made the strain chitosan-deficient similar to the *cda1Δ2Δ3Δ* of *C. neoformans*. The majority of the *in vitro* phenotypes of the *cda1Δ2Δ3Δ* strain of R265 were similar to the *cda1Δ2Δ3Δ* strain made in *C. neoformans*. However, *cda1Δ2Δ3Δ* of *C. gattii* R265 grew better in YPD (with markedly less morphological abnormalities) than the *cda1Δ2Δ3Δ* of *C. neoformans* (data not shown). *C. neoformans cda1Δ2Δ3Δ* exhibited a leaky melanin phenotype when incubated in L-DOPA medium (28). However, the *cda1Δ2Δ3Δ* οf R265 did not show this phenotype (Supplementary Fig. 6) suggesting that there are differences in the cell wall architecture between *cda1Δ2Δ3Δ* cells of *C. neoformans* and *C. gattii*. In *C. neoformans* we have shown that *CDA1* is specifically upregulated during the growth of yeast cells in the infected lung. Recent whole genome transcriptome studies involving the four lineages of *C. gattii* have shown that *C. gattii* R265 displays a distinct differential gene expression profile compared to the rest of the species under *in vitro* and *ex vivo* (incubation of the yeast cells with bone marrow derived macrophages) growth conditions (36). Significant differential expression of genes involved in capsule biosynthesis, cell wall remodeling and genes involved in other virulence related traits have been shown in two separate studies of *C. gattii* R265 (36, 37). Specifically, in *C. gattii* R265 the *CDA1* and *CDA2* genes were found to be significantly down regulated when the cells were incubated with bone marrow derived macrophages (36). Taken together, these results and our data demonstrating the essential role of Cda3 in *C. gattii* R265 virulence suggest a transcriptional regulation of chitosan biosynthesis during mammalian infection.

We have previously shown that vaccination with chitosan-deficient *cda1Δ2Δ3Δ* strain of *C. neoformans* at the optimal concentration confers robust protective immunity to subsequent challenge infection with wild-type virulent KN99 (29). Even though deletion of all three CDA genes of *C. gattii* produced a chitosan-deficient strain, vaccination with either the live strain or a heat killed preparation of it induced only partial protection to subsequent infection with R265. This may be due to the fact that R265 has the inherent ability to dampen host-induced immune responses (22–25).

In summary, the hypervirulent *C. gattii* strain R265 has evolved with a distinct transcriptional profile with altered expression of components of chitin deacetylation that may have enabled it to adapt more efficiently to various environmental conditions with ramifications on its virulence. Of the several reported differences in the virulence related traits between *C. neoformans* and *C. gattii* R265, the difference in the regulation of chitosan biosynthesis as revealed from our studies may significantly contribute to the mechanisms of its unique and distinct nature of pathogenesis since chitosan is one of the macromolecules that resides at the host pathogen interface.

## Materials and methods

### Fungal strains and media

R265, *C. gattii* strain of VGII subtype linked to the 1999 British Columbia outbreak (11), was used as the wild-type strain and as progenitor of mutant strains. This strain was kindly provided by Joseph Heitman (Duke University Medical Center, NC). All the strains used in this study are listed in Table 1. KN99α, a strain of *C. neoformans*, was used as the wild -type strain for serotype A (31). Strains were grown on YPD (1% yeast extract, 2% bacto-peptone, and 2% dextrose). Solid media contained 2% bacto-agar. Selective YPD media contained 100 μg/mL nourseothricin (NAT) (Werner BioAgents, Germany) and/or 200 μg/mL G418 (Geneticin, Gibco Life technologies, USA). RPMI 1640 (Corning 10-040 CM) contained 10% fetal bovine serum (FBS, Gibco-Thermo Fisher Technologies, # 26140).

### Generation of deletion constructs of *C. gattii*

Gene-specific deletion constructs of the chitin deacetylases were generated using overlap PCR gene technology described previously (38, 39) and included either the hygromycin resistance, geneticin resistance cassette (40) or nourseothricin resistance cassette (41). The primers used to disrupt the genes are shown in Table S1. The Cda1 deletion cassette contained the nourseothricin resistance cassette resulting in a 1,539 bp replacement of the genomic sequence between regions of primers 3-Cda1 and 6-Cda1 shown in upper case in Table S1. The Cda2 deletion cassette contained the hygromycin resistance cassette resulting in a 1,587 bp replacement of the genomic sequence and the Cda3 deletion cassette contained the hygromycin resistance cassette resulting in a 1,494 bp replacement of the genomic sequence. Constructs were introduced into the R265 strain using biolistic techniques (42).

### Transformation and characterization of *C. gattii* mutants

Recipient strains of *C. gattii* were transformed biolistically following the protocol described earlier (40, 42). Drug resistant transformants that formed colonies in 3-5 days were passaged four times in liquid YPD medium before reselection of drug resistance on agar. Transformants were further screened by diagnostic PCR of their genomic DNA using primers at the 5’ and 3’ junction of the integration site of the transforming DNA. Southern blot hybridizations were done to verify the absence of random DNA integrations, as described previously employing DIG labelled DNA probes (43, 44).

### Cellular chitosan measurement

As previously described, MBTH (3-methyl-2-benzothiazolinone hydrazone) based chemical method was used to determine the chitin and chitosan content of *C. gattii* or *C. neoformans* (31). In brief, cells were collected after growing them in appropriate media and growth conditions by centrifugation. Cell pellets were washed two times with PBS, pH 7.4 and lyophilized. The dried samples were resuspended in water first before adding KOH to a final concentration of 6% KOH (w/v). The alkali suspended material was incubated at 80°C for 30 min with vortexing in between to eliminate non-specific MBTH reactive molecules from the cells. Alkali treated material was then washed several times with PBS, pH 7.4 to make sure that the pH of the cell suspension was brought back to neutral pH. In the case of the cells grown in RPMI, alkali treated material was sonicated as described previously to generate a uniform suspension (30). Finally, the cell material was resuspended in PBS, pH 7.4 to a concentration of 10 mg/mL in PBS (by dry weight) and a 0.1 mL aliquot of each sample was used in the MBTH assay (45).

### Virulence and fungal burden assays

*C. gattii* strains were grown at 30°C, 300 rpm for 48 hours in 50 mL YPD. The cells were centrifuged, washed in endotoxin-free 1x PBS and suspended in 5 mL of the same. The cells were counted with a haemocytometer and diluted to 2x 10^6^ cells/mL. CBA/J female mice (Jackson Laboratories) were anaesthetized with an intraperitoneal injection (200 µL) of ketamine (8 mg/mL)/dexmedetomidine (0.05 mg/mL) mixture, which was reversed by an intraperitoneal injection of (200 µL) of antipamezole (0.25mg/mL). Mice were allowed to inhale 1×10^5^ cells in 50 μL, which were dripped into the nares. For virulence assays, mice were weighed before and during the course of infection. Mice were euthanized by CO_2_ asphyxiation if they reached 80% of their original body weight. At this point, the mice appeared morbidly ill displaying a ruffled coat, lethargy, a hunched posture, unstable gait and loss of appetite. For the determination of CFUs, lung or brain from each mouse was placed in 2.0 mL of 1x PBS (pH 7), homogenized, serially diluted, plated onto YPD agar supplemented with 100 µg/mL streptomycin and 100 µg/mL ampicillin, and incubated for 2 days at 30°C. Total CFUs per organ were calculated. The infection protocol was reviewed and approved by the Washington University School of Medicine Animal Care and Use Committee (IACUC).

### Evaluation of *C. gattii* to stress under in vitro conditions

Solid YPD medium was made with the desired amount either SDS, NaCl, calcofluor white, or Congo red. For plating, wild-type and mutant strains were grown in liquid YPD for 24 hours at 30°C. Cells were diluted to OD_650_=1.0 and 10-fold serial dilutions were made. Five microliters of each dilution were spotted on the plate and the plates were incubated for 2-3 days at appropriate temperatures and photographed. Eosin Y staining was carried out as described earlier (29).

### Analysis of melanin production

Strains were grown overnight in 2 mL YPD medium at 30°C with shaking to saturation. Cells were collected, washed in 1xPBS. Then, 1×10^8^ (5×10^7^ /ml) of each mutant was added to 4 mL of glucose-free asparagine media (1 g/liter L-asparagine, 0.5g/liter MgSO_4_ 7H_2_O, 3 g/liter KH_2_PO_4_, and 1mg/liter thiamine, plus 1 mM L-3,4-dihydroxyphenylalanine (L-DOPA) for 7 days at 300 RPM and 30°C in the dark. Samples were then spun down at >600 x g for 10 minutes. The cells’ ability to produce pigment was assessed visually.

### Capsule analysis

Cells were grown in YPD at 30°C. After 48 hrs for growth, cells were collected, washed once with PBS and were inoculated to RPMI medium containing 10% fetal bovine serum (FBS) at a concentration of 500 cells/µL and incubated for five days at 37°C in the presence of 5% CO_2_. The capsule-induced strains were resuspended in a 1:4 India ink:H_2_O solution and photographed on an Olympus BX61 microscope. Capsule diameter was measured and averaged for a minimum of 100 cells per strain using SlideBook 5.0 (Intelligent Imaging Innovations, Inc. CO, USA).

### Statistics

Data were analyzed using GraphPad Prism, version 7.0 (GraphPad Software, Inc., La Jolla, CA). The unpaired two-tailed t test with Welch’s correction was used for comparisons of two groups. The one-way analysis of variance (ANOVA) with the Dunnett’s multiple-correction test was used to compare more than two groups. Kaplan-Meier survival curves were compared using the Mantel-Cox log rank test.

## Funding

This work was supported by NIH grants R01AI072195 to JKL; R01AI125045 to JKL, CAS and SML and R01AI025780 to CAS and SML.

## Supplementary Materials

**Figure S1:**
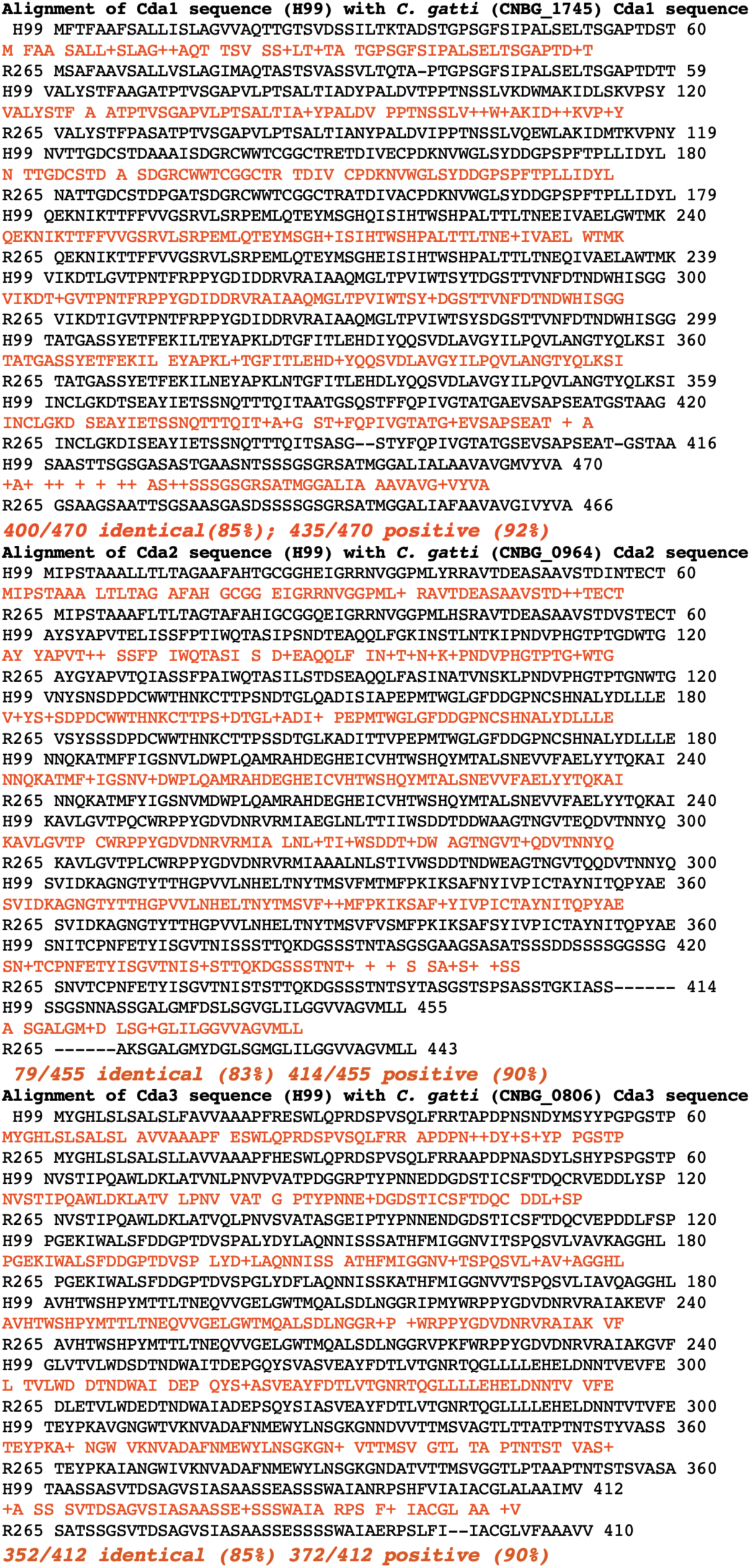
Pair-wise alignment of CDA protein sequences of *C. neoformans* and *C. gattii* R265.

**Figure S2.**
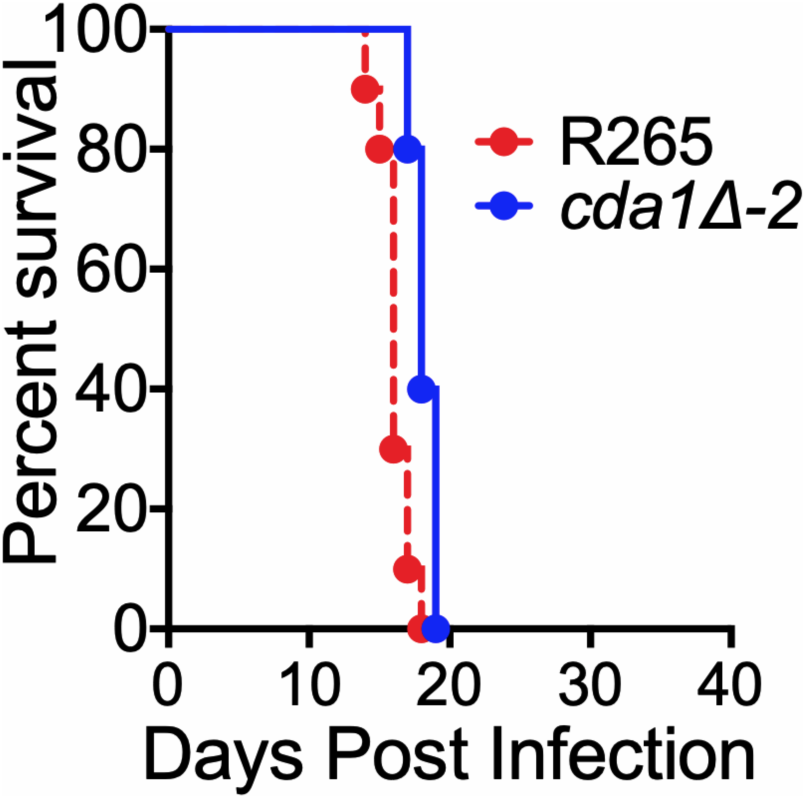
Deletion of *CDA1* in R265 does not affect fungal virulence. Mice were inoculated with 50,000 CFU of either wild-type R265 or the second isolate of *cda1Δ*. Survival of the animals was recorded as mortality. Mice that lost 20% of the body weight at the time of inoculation were considered ill and sacrificed.

**Figure S3.**
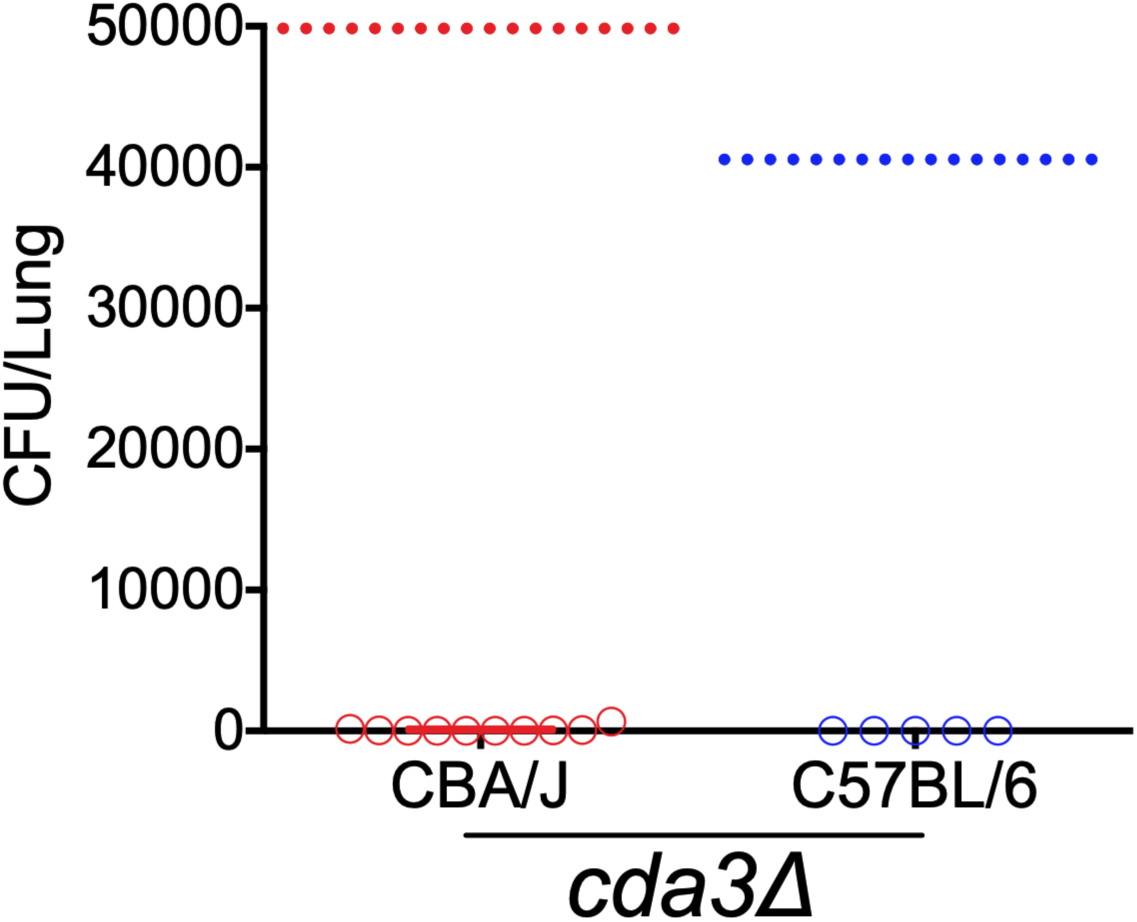
Fungal burden in the lungs of the mice at the end point of the survival experiment. Fungal burden in the lungs of the mice at the end point of the survival experiment for both CBA/J and C57BL/6. The dashed line indicates the CFU of the initial inoculum for each mouse strain.

**Figure S4.**
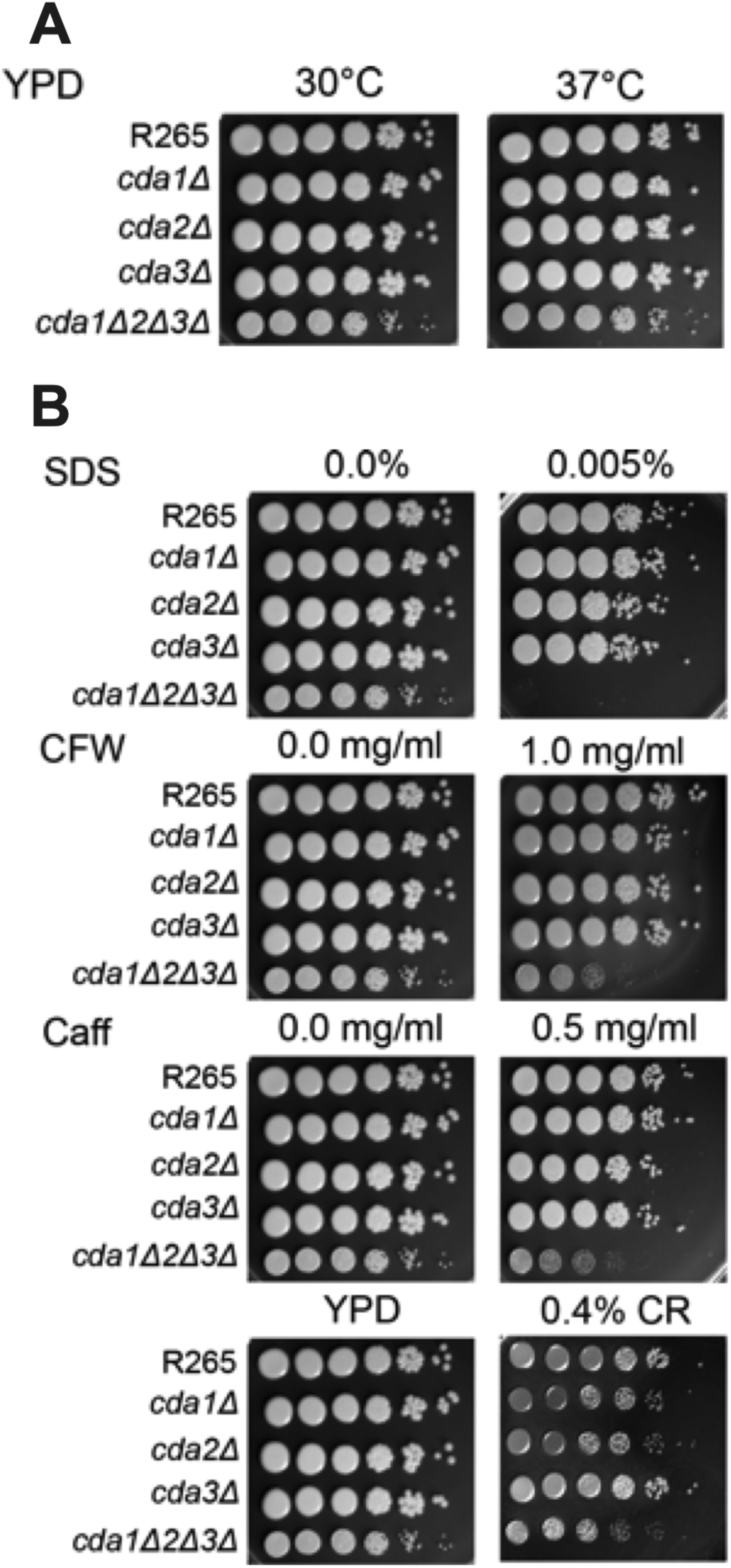
Sensitivity of chitin deacetylase mutants to cell wall inhibitors and temperature. Cultures were grown overnight in YPD then diluted to an OD_650_ of 1.0. Tenfold serial dilutions were made in PBS and 5µl of each was plated. The plates were grown for 5 days at 30°C for the inhibitor plates and at the indicated temperature for all others. The wild-type (R265) and deletion stains are labelled on the left and the conditions noted at the top. (**A**). Sensitivity of mutants to temperature. (**B**). Sensitivity to cell wall inhibitors. CFW (calcofluor white), SDS (sodium dodecyl sulphate), Caff (caffeine) and CR (Congo red).

**Figure S5.**
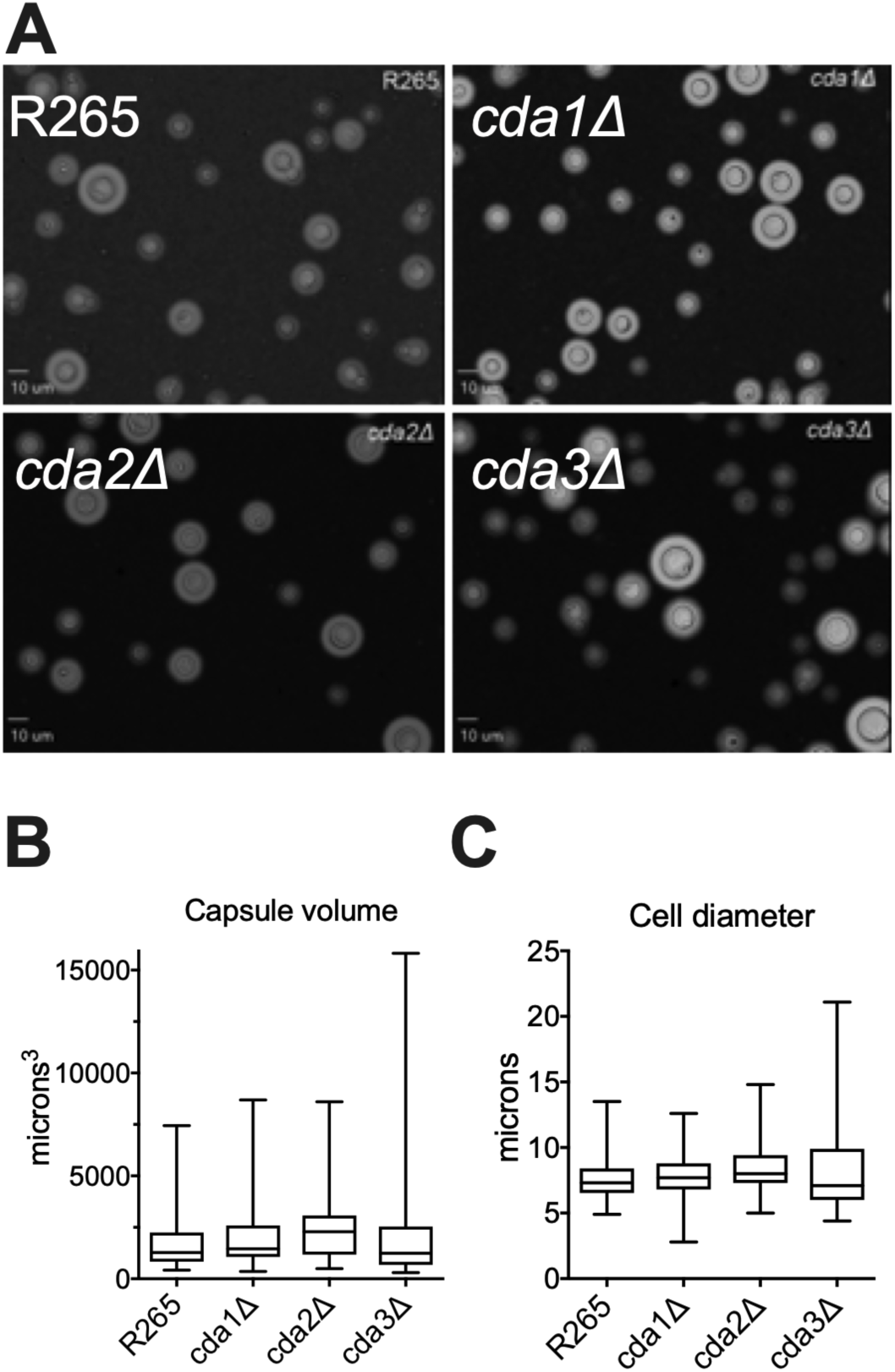
*C. gattii* R265 *cda1Δ*, *cda2Δ* or *cda3Δ* mutants display normal levels of capsule under capsule inducing conditions. Cells were incubated under capsule-inducing conditions for 5 days. Capsule size was assessed by staining with India ink and visualizing the zone of exclusion at a magnification of ×60.

**Figure S6.**
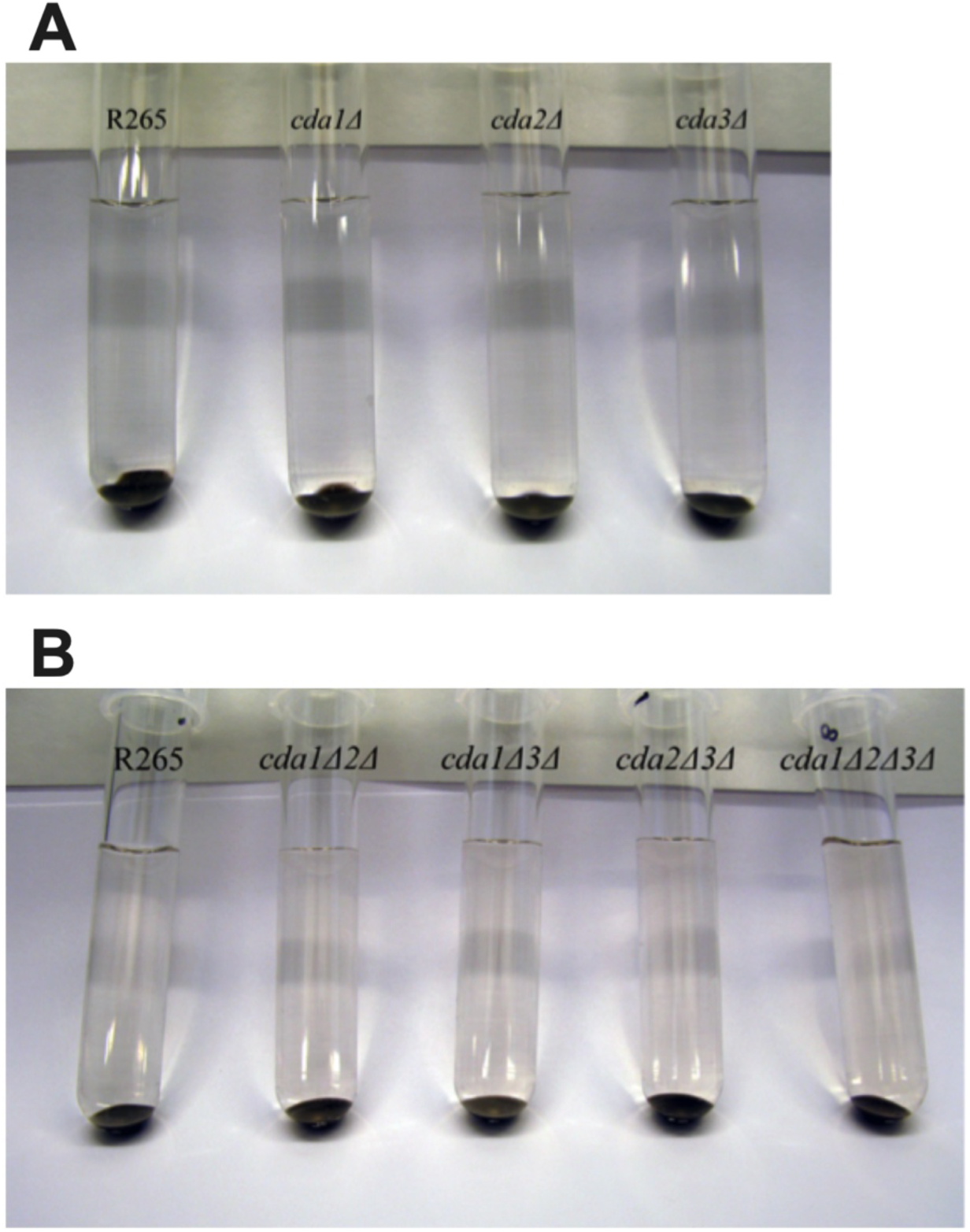
Melanin phenotype of chitin deacetylase mutants of *C. gattii*. Chitin deacetylase mutants displayed no difference in melanin production compared with wild type (R265). Strains were grown overnight in 2 mL YPD medium at 30°C with shaking to saturation. Cells were collected, washed in 1xPBS. Then, 1×10^8^ (5×10^7^ /ml) of each mutant was added to 4 mL of Asn+L-DOPA for 7 days at 300 RPM and 30°C in the dark. Samples were then spun down and photographed. Black pellets indicate the presence of melanin in the strain. The clear supernatants indicate that there was not a “leaky melanin” phenotype (27, 28).

**Figure S7.**
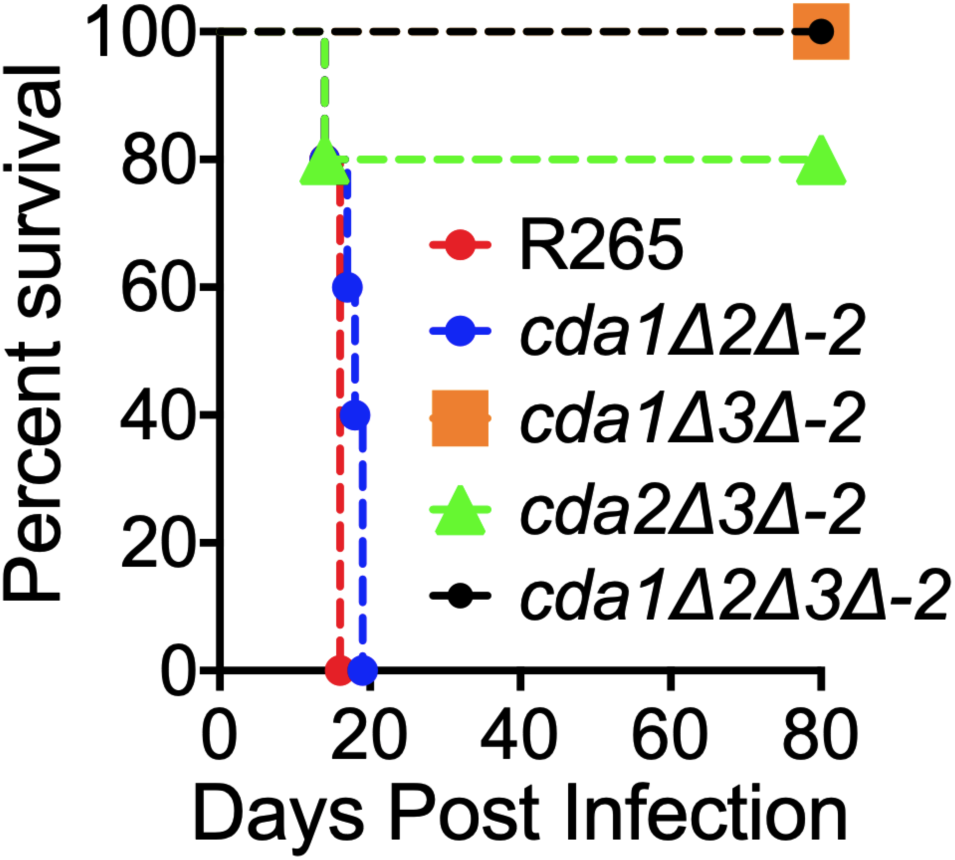
Deletion of *C. gattii CDA3* in combination with any of the other two CDAs results in severe attenuation of virulence in mouse model of infection. CBA/J mice (6-8 weeks old, female) were infected intranasally with 10^5^ CFU of second independent isolate of each strain. Survival of the animals was recorded as mortality of mice for 80 days PI. Mice that lost 20% of the body weight at the time of inoculation were considered ill and sacrificed. Data is representative of two independent experiments with five animals for each strain.

**Figure S8.**
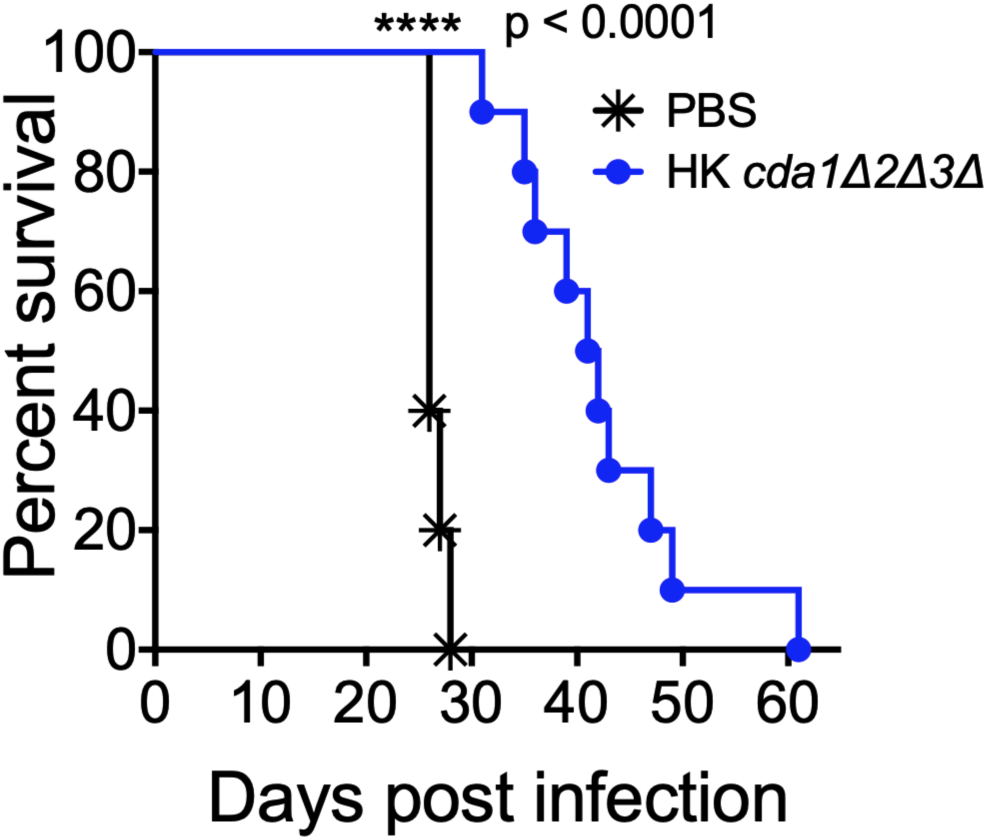
Vaccination with a heat-killed preparation of *cda1Δ2Δ3Δ* of *C. gattii* confers attenuated protection to a subsequent challenge infection with virulent R265. Mice (C57BL/6) were immunized with a HK preparation of *cda1Δ2Δ3Δ* strain with a dose equivalent to 10^7^ CFU. Control mice were inoculated with PBS. After 40 days, mice were challenged with 10,000 CFU of wild-type R265 cells. Survival of the animals was recorded as described above.

**Table S1.**
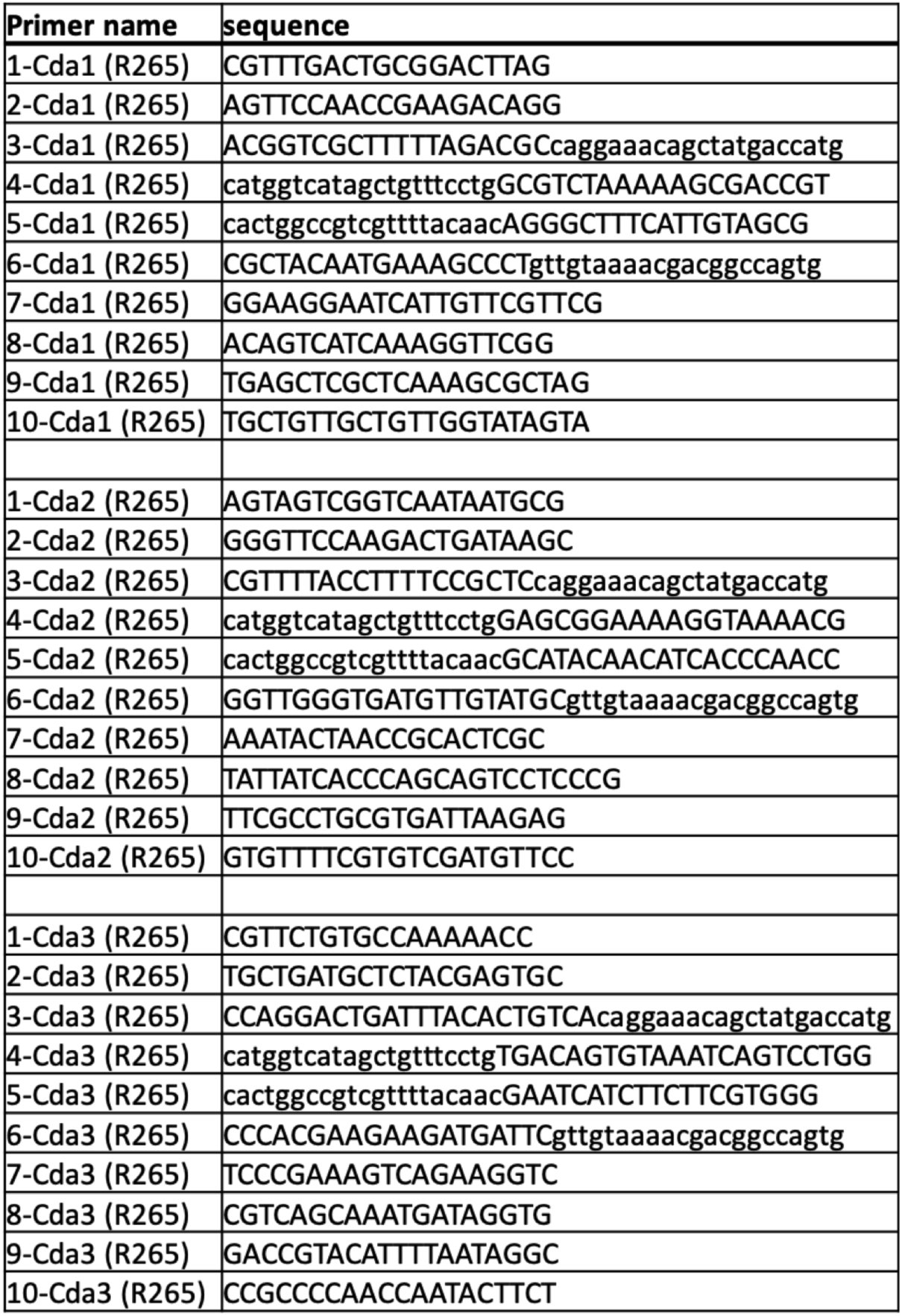
Primers used in this study.

